# Independent evolution of harmala alkaloids in medicinal Syrian rue and ayahuasca plants

**DOI:** 10.64898/2026.10.24.761205

**Authors:** Fabiola Muro-Villanueva, Ye Won Ham, Justin Williams, Yujing Yan, Danielle E. Khost, Juan A. Orozco, Timothy B. Sackton, Charles C. Davis, Ryan S. Nett

## Abstract

Plants from distantly related lineages can produce identical neuroactive molecules, many of which have been independently adopted across human societies for their medicinal and cognition-altering effects. This phenomenon is exemplified by the traditional use of Syrian rue (*Peganum harmala*) and the ayahuasca vine (*Banisteriopsis caapi*) for medicine and ritual. Despite diverging over 100 million years ago and inhabiting different continents and climates, both plants synthesize harmala alkaloids (HAs), which inhibit human monoamine oxidase and exhibit diverse neuroactive effects. Here, we combine gene expression analyses, genomics, and enzyme characterization to elucidate HA biosynthesis for both Syrian rue and ayahuasca vine. We reveal multiple layers of biochemical and genetic convergence in these pathways, including independent duplication and neofunctionalization of an ancestral enzyme from serotonin biosynthesis, and lineage-specific repurposing of primary metabolic genes to generate a reactive precursor for alkaloid scaffold formation. Our study resolves biosynthetic routes to HAs in two medicinal plants and demonstrates convergent evolution of specialized metabolism at the levels of enzymes, pathways, and genomes.

## INTRODUCTION

Plants produce thousands of natural products that influence animal behavior^1^. Yet despite this chemical diversity, distantly-related plant lineages have repeatedly evolved the capacity to produce the same neuroactive molecules. Many of these convergently-evolved molecules, including caffeine, cannabinoids, and tropane alkaloids^2–4^, have been important throughout human history as regulators of mood, behavior, and cognition^5^. Indeed, plants that produce similar neuroactive chemistry have also been independently adopted by different human cultures for similar purposes. A prime example of this convergent adoption of neuroactive plants is the historical use of Syrian rue (*Peganum harmala* L.) and ayahuasca vine (*Banisteriopsis caapi* (Griseb.) C. V. Morton)) as both medicines and entheogens. These plants belong to plant orders that diverged more than 100 million years ago and inhabit very different geographic and ecological settings (**Fig. 1**): *P. harmala* across Eurasia and North Africa, and *B. caapi* in Amazonia. Despite these differences, both plants produce harmala alkaloids (HAs)^6^, a class of β-carboline-related molecules that inhibit human monoamine oxidase (MAO)^7^, a major pharmacological target of antidepressants^8^. The MAO inhibitory activity of HAs is also central to the traditional use of *B. caapi* in ayahuasca decoctions, which are multi-plant preparations that have been used by Indigenous Amazonian groups for at least 1000 years for spiritual and medicinal purposes^9^. In these preparations, HAs from *B. caapi* inhibit human MAO and thereby reduce the breakdown of ingested dimethyltryptamine (DMT), a psychedelic compound that is supplied by other plant ingredients, including *Psychotria viridis*^10^. Similarly, *P. harmala* has been used for nearly 3000 years in Eurasia and North Africa for its consciousness-altering properties, to treat a variety of medical ailments, and as an insect repellent^11,12^. These diverse bioactive effects are thought to be linked to the high levels of HAs present in this plant.

**Figure 1.**
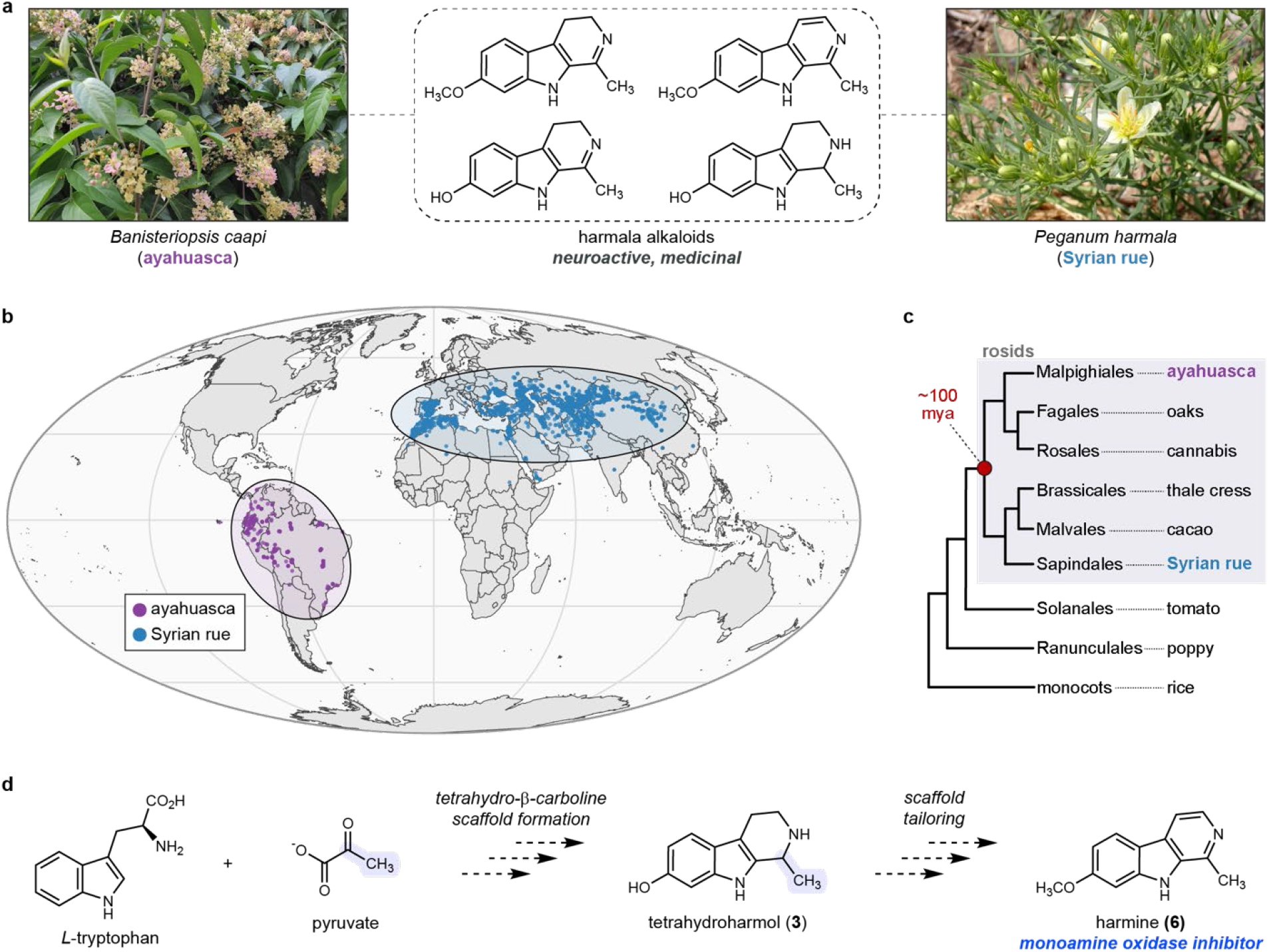
Distinct origins and geography of plants that produce harmala alkaloids. **a)** *Banisteriopsis caapi* (ayahuasca) and *Peganum harmala* (Syrian rue) make identical, medicinal harmala alkaloids. **b)** Distinct endemic geographic distributions of ayahuasca and Syrian rue. **c)** Ayahuasca and Syrian rue belong to plant families that diverged around 100 million years ago (mya). Shown is a representative phylogeny of major angiosperm taxa. **d)** Proposal for harmine (**6**) biosynthesis based on prior isotope tracer studies and chemical logic.

More than 60 HAs have been identified across the plant kingdom^6^. Although their biological functions remain poorly understood, HAs act as insecticides^12^ and can inhibit plant growth^13^, suggesting diverse roles in plant ecology. Due to their structural similarity to neurotransmitters and broader neuroactive effects, HAs have been explored as promising treatments for numerous neurological disorders, including depression and Alzheimer’s disease^14^. Additionally, recent work has shown that harmine, the most abundant HA in *P. harmala* and *B. caapi*, has promise as a therapeutic for diabetes^15,16^. Despite this ecological, cultural, and biomedical importance, little is known about how plants synthesize HAs, including whether their biosynthetic pathways evolved independently in different plant lineages. Previous isotope-labeling in *P. harmala* established that harmine (**6**) is derived from *L*-tryptophan and pyruvate in this species (**Fig. 1**)^17,18^. However, no HA biosynthetic genes have been identified from either *P. harmala* or *B. caapi*. HA biosynthesis therefore provides an opportunity to investigate how distantly related plants evolve the same neuroactive chemistry.

Here, we hypothesized that resolving HA biosynthesis in *P. harmala* and *B. caapi* could illuminate its evolutionary origin across biological scales, from metabolites and biochemical pathways to enzymes and genomes. More broadly, these plants provide a powerful model for integrating ethnopharmacology, biochemistry, and molecular biology to understand the complex relationships between plants and humans^19^. By combining analytical chemistry, comparative transcriptomics and genomics, and functional enzyme characterization, we reconstructed HA biosynthesis for both *P. harmala* and *B. caapi*. In addition to providing pathways to HAs in two medicinal plants, our results demonstrate how independent evolutionary histories can generate the same chemical outcome via both shared and lineage-specific innovations.

## RESULTS

### Discovery of a harmine biosynthetic pathway in *Peganum harmala*

The active production of plant specialized metabolites typically correlates with biosynthetic gene expression, and this can be leveraged for biosynthetic pathway elucidation^20–22^. Thus, we first focused on identifying where HAs are actively biosynthesized in *P. harmala* seedlings (**Fig. 2a**). Using liquid chromatography-mass spectrometry (LC-MS) we observed that the major HAs in *P. harmala* (harmalol (**4**), harmaline (**5**), and harmine (**6**)) were found throughout the seedling, but were most abundant in the roots (**SI Fig. 1**). To measure active biosynthesis of harmine (**6**), we fed *P. harmala* seedlings with deuterated *L*-tryptophan (1 mM D_5_-Trp). The labeled substrate reached all tissues (**SI Fig. 2a, 2b**), but the highest incorporation into HAs occurred in roots (**Fig. 2b, SI Fig. 2c**), supporting this as the site of HA biosynthesis in seedlings.

**Figure 2.**
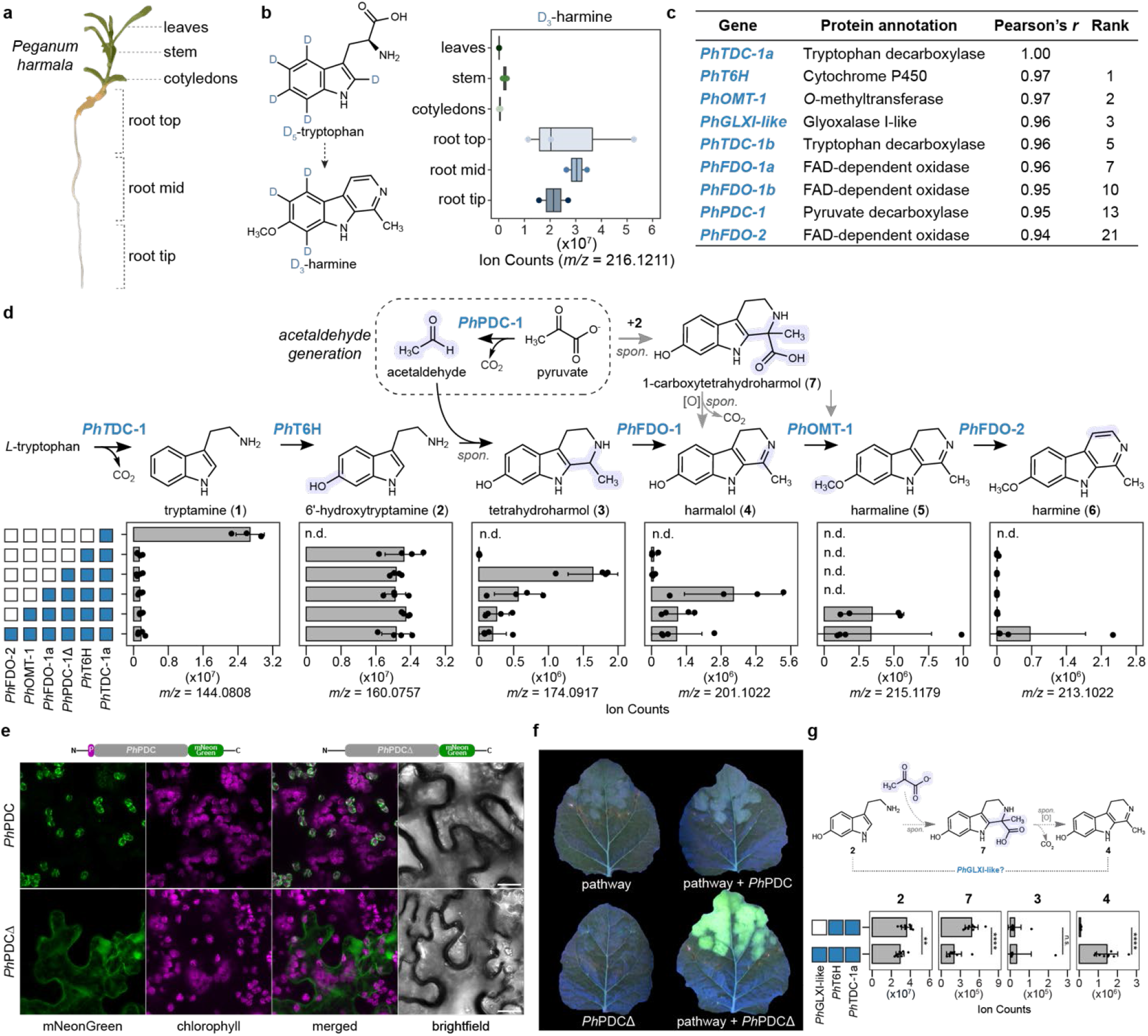
Total biosynthesis of harmine in Syrian rue. **a)** Syrian rue (*Peganum harmala*) seedling. Annotated tissues were used for isotope labeling studies and RNA-sequencing. **b)** Isotope labeling with deuterated tryptophan (D_5_-tryptophan) defined tissues with active harmine (**6**) biosynthesis. Shown are quantified extracted ion chromatograms (EICs) for D_3_-harmine (**6**) in different *P. harmala* tissues. **c)** Gene co-expression analysis identifies gene candidates with strong co-expression to a tryptophan decarboxylase gene (*PhTDC-1a*) from *P. harmala*. **d)** Stepwise reconstitution of harmine (**6**) biosynthesis in Syrian rue using transient expression in *N. benthamiana*. Shown are quantified EICs for the mass ion (*m/z*) of each intermediate. Dots represent biological replicates, with bar graphs depicting the mean ± s.d. (*n* = 4). Blue boxes to the left of bar graphs indicate inclusion of a gene in transient expression experiments, while white boxes indicate omission of a gene. spon. = spontaneous., n.d. = not detected. **e)** Confocal microscopy images visualizing the localization of mNeonGreen-fused *Ph*PDC or *Ph*PDCΔ (N-terminal signal peptide removed) proteins during transient expression in *N. benthamiana* cells. Scale bar = 20 µm. **f)** Harmala alkaloid production can be observed via UV fluorescence when *Ph*PDCΔ is included with the rest of the pathway genes. **g)** Transient co-expression of *Ph*GLXI-like with *Ph*TDC-1a and *Ph*T6H in *N. benthamiana* affects the accumulation of harmala alkaloid pathway intermediates. Shown are quantified EICs for the mass ion (*m/z*) of each relevant intermediate listed above individual bar graphs. Dots represent biological replicates, with bar graphs depicting the mean ± s.d. (*n* = 9). For 1-carboxytetrahydroharmol (**7**), an in-source fragment was used for quantification ([M-CH_2_O_2_+H]^+^ = *m/z* 201.1022).

We next used long read RNA-sequencing (RNA-seq) to generate a reference transcriptome, and profiled gene expression with short read RNA-seq to identify candidate biosynthetic genes with enriched expression in the roots. The sequenced samples included multiple tissue types with high and low biosynthetic activity (**Fig. 2a**), as this has proven to be a powerful strategy for identifying co-expressed biosynthetic genes^20–22^. Previous work demonstrated that tryptamine (**1**), the product of *L*-tryptophan decarboxylation, is incorporated into HAs^17^. This suggested that biosynthesis is initiated via tryptophan decarboxylase (TDC), a member of the pyridoxal phosphate (PLP)-dependent decarboxylase family that is known to catalyze this reaction in plants^23^. We looked for PLP-dependent decarboxylases with enriched expression in the roots and filtered for candidates that had conserved active site residues characteristic of TDCs^23^. Among the 45 PLP-dependent decarboxylase candidates identified in the transcriptome, only two, *Ph*TDC-1a and *Ph*TDC-1b (98% amino acid identity), met these criteria (**SI Fig. 3a**). *Ph*TDC-1a and *Ph*TDC-1b were characterized in *Nicotiana benthamiana* using Agrobacterium-mediated transient expression^24^, and both enzymes decarboxylated endogenous *L*-tryptophan from *N. benthamiana* to form tryptamine (**1**) (**SI Fig. 3b**).

**Figure 3.**
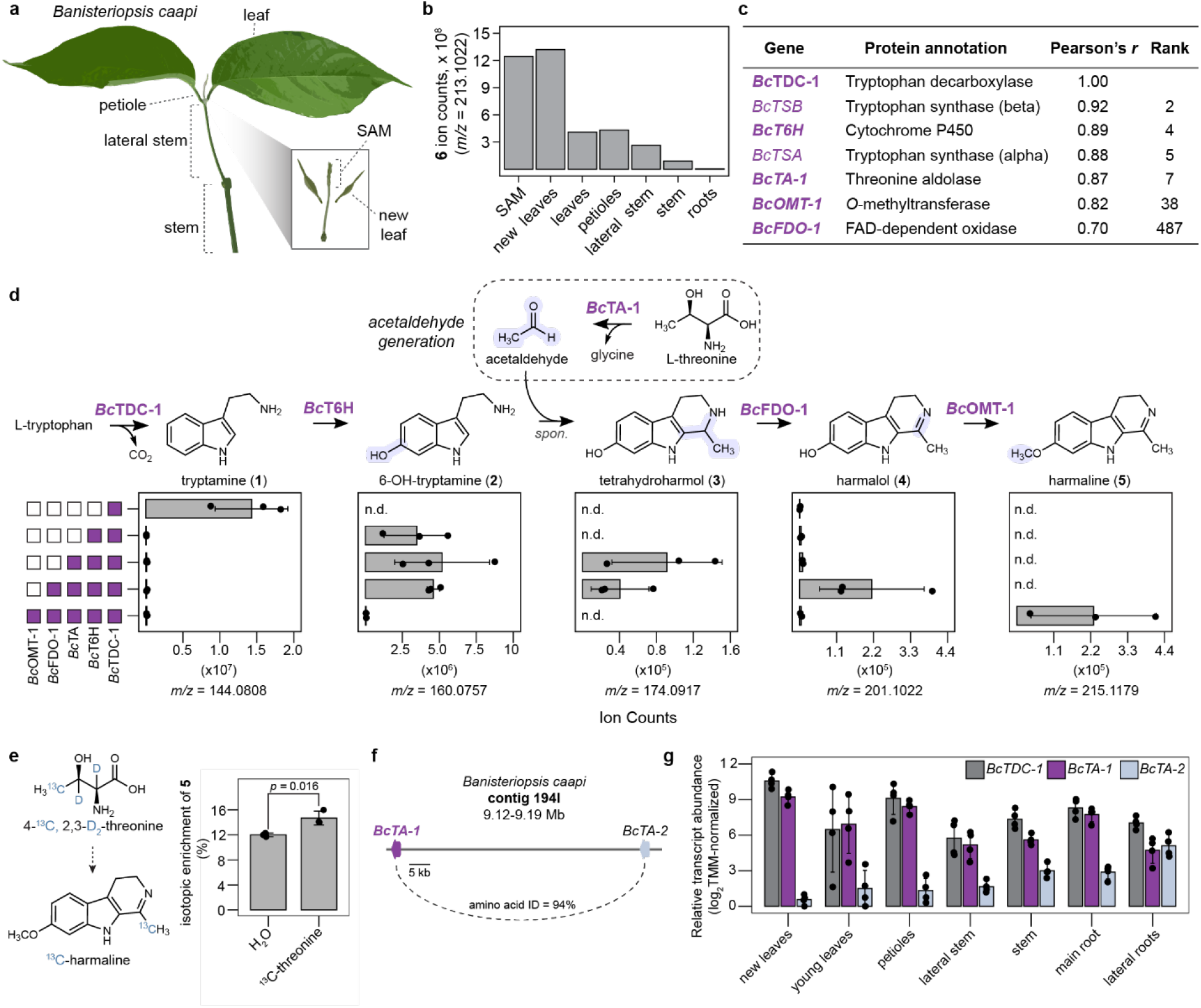
A distinct harmala alkaloid biosynthetic pathway in ayahuasca. **a)** Schematic of ayahuasca (*Banisteriopsis caapi*) aerial tissues that were used for harmala alkaloid profiling and RNA-sequencing. **b)** Distribution of harmine (**6**) in ayahuasca tissues, quantified here by ion abundance normalized to tissue fresh weight. **c)** Co-expression analysis using Bc*TDC* as a bait gene identifies candidate enzymes for harmala alkaloid biosynthesis. Genes that are bolded were functionally characterized here. **d)** Step-by-step reconstitution of ayahuasca harmala alkaloid biosynthesis using transient expression in *N. benthamiana*. Shown are quantified EICs for the mass ion (*m/z*) of each intermediate. Dots represent biological replicates, with bar graphs depicting the mean ± s.d. (*n* = 3). Purple boxes to the left of bar graphs indicate inclusion of a gene in transient expression experiments, while white boxes indicate omission of a gene. spon. = spontaneous., n.d. = not detected **e)** Labeled threonine is incorporated into harmaline (**5**) in ayahuasca. Shown is the labeling percentage of threonine calculated by dividing labeled threonine (^13^C, D_2_-threonine) by the sum of labeled and unlabeled threonine in ayahuasca plants (*n* = 3) treated with H_2_O (control) or labeled threonine. **g)** Tandem duplication of the threonine aldolase (TA) gene in the ayahuasca genome. **g)** Tissue-specific expression of *BcTA-1* and *BcTA-2* compared to *BcTDC* in ayahuasca tissues that were profiled via RNA-seq. Dots represent biological replicates, with bar graphs depicting the mean ± s.d. (*n* = 4).

We then used *Ph*TDC-1a as a bait to identify additional genes with similar expression patterns that may act in HA biosynthesis (**Supplementary Data 1**). We initially focused on the downstream tailoring of the tetrahydro-β-carboline scaffold, which would require oxidase (e.g., cytochrome P450 monooxygenase or flavin-dependent oxidase) and methyltransferase enzymes. Strikingly, of all 73,698 transcripts from our *P. harmala* transcriptome, a cytochrome P450 (CYP), a putative *O*-methyltransferase (OMT), and three flavin-dependent oxidase (FDO) genes were within the top 23 co-expressed genes with *Ph*TDC-1a (**Fig. 2c, Supplementary Data 1**).

By transiently co-expressing the CYP (*Ph*T6H; CYP71BG47) with *Ph*TDC-1a in *N. benthamiana*, we found that it acts as tryptamine 6ʹ-hydroxylase (T6H) to convert tryptamine (**1**) into 6ʹ-hydroxytryptamine (**2**) (**Fig. 2d**, **SI Fig. 4**). Downstream oxidase and methyltransferase reactions were tested by co-infiltrating HAs as substrates with our transiently expressed OMT and FDO gene candidates. The putative OMT (*Ph*OMT-1) had *O*-methyltransferase activity and catalyzed the following reactions: tetrahydroharmol (**3**) to tetrahydroharmine (**8**), harmalol (**4**) to harmaline (**5**), and harmol (**9**) to harmine (**6**). (**Fig. 2d**, **SI Fig. 4, 5**). Moreover, two FDO enzymes (*Ph*FDO-1a and *Ph*FDO-1b; 82% amino acid identity) oxidized tetrahydro-β-carbolines (tetrahydroharmol (**3**) and tetrahydroharmine (**8**)) into the corresponding dihydro-β-carboline products (harmalol (**4**) and harmaline (**5**)) (**Fig. 2d**, **SI Fig. 4, 6**). The third FDO (*Ph*FDO-2) oxidized harmaline (**5**) to yield harmine (**6**) (**Fig. 2d**, **SI Fig. 4, 6**). The relatively high amino acid identity (80%) between *Ph*FDO-1a/*Ph*FDO-1b and *Ph*FDO-2 suggested that *Ph*FDO-2 may have arisen through the duplication and neofunctionalization of a *Ph*FDO-1 homolog. Indeed, in addition to oxidizing harmalol (**4**) and harmaline (**5**), *Ph*FDO-2 retains low activity to oxidize tetrahydroharmine (**8**) into harmaline (**5**) (**SI Fig. 6**). Furthermore, we mined the published genome of *P. multisectum*^25^, a relative of *P. harmala* that also produces HAs, and found that the three FDO gene orthologs are adjacent to each other within a 30 kilobase (kb) region of chromosome 6 (**SI Fig. 7**). Altogether, these data support a duplication and neofunctionalization event in *Peganum* that gave rise to the FDOs involved in HA biosynthesis. Our results are also consistent with the recent report of orthologous TDC (98% amino acid identity) and OMT (96.1% amino acid identity) enzymes from *P. multisectum*^26^.

**Figure 4.**
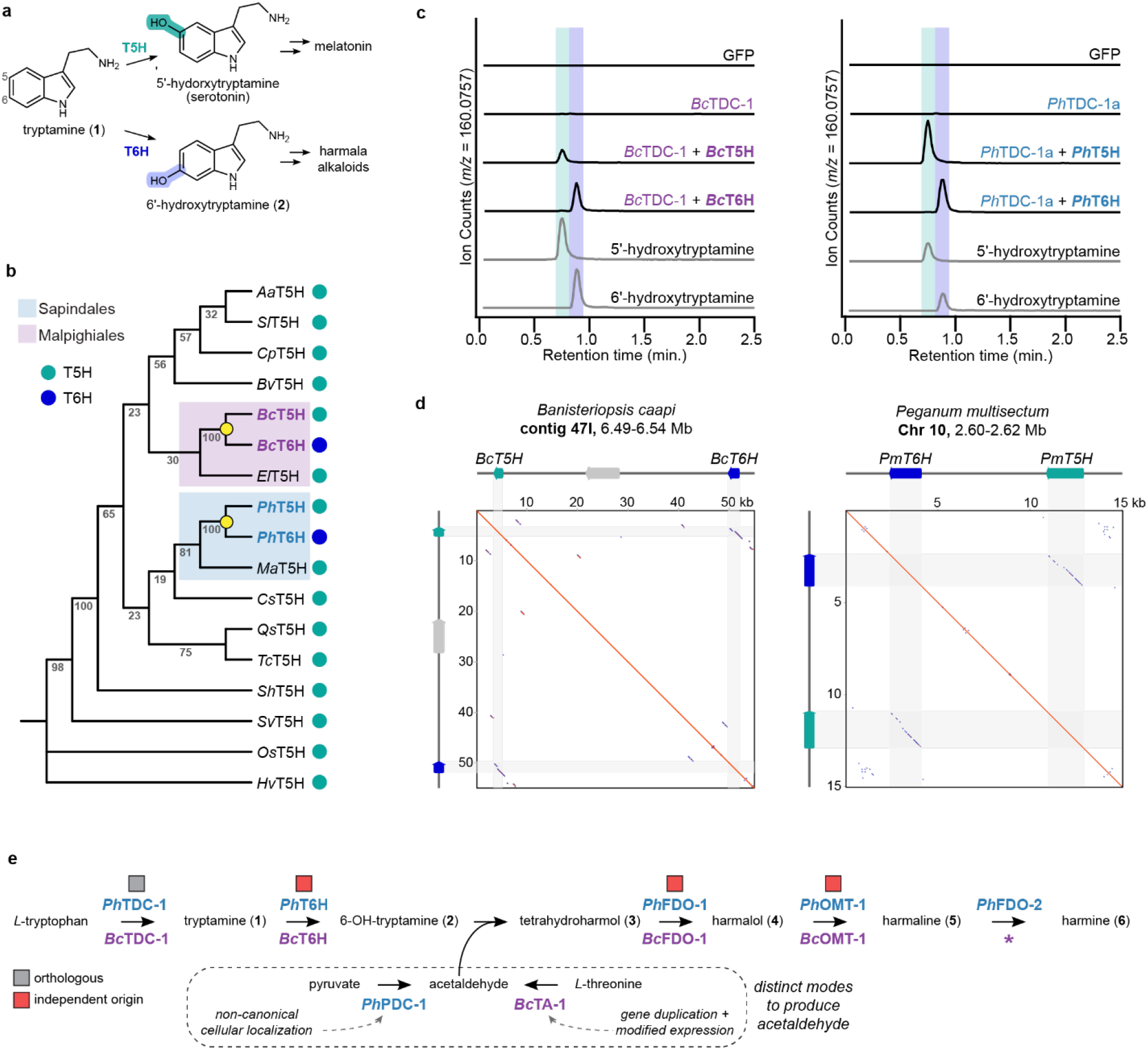
Independent origins of harmala alkaloid biosynthesis in Syrian rue and ayahuasca. **a)** Comparison of tryptamine 5-hydroxylase (T5H) and tryptamine 6-hydroxylase (T6H) enzyme activities. **b**) Simplified phylogenetic tree of *Bc*T6H and *Ph*T6H compared to verified T5H genes from across the plant kingdom. Yellow dots indicate the ancestral nodes for T6H genes from Syrian rue and ayahuasca. Branch lengths are not representative of evolutionary distance, see SI Fig. 20 for complete tree. **c)** Characterization of T5H and T6H enzymes from *B. caapi* (left) and *P. harmala* (right) by LC-MS. Shown are EICs for 5ʹ-hydroxytryptamine and 6ʹ-hydroxytryptamine ([M-NH_3_+H]^+^ = *m/z* 160.0757) **d)** Duplication and neofunctionalization of the T5H gene in the *B. caapi* (left) and *P. harmala* (right) genomes with corresponding dot plots to demonstrate sequence similarity between T5H and T6H loci. **e)** Comparison of harmala alkaloid biosynthetic pathways in ayahuasca and Syrian rue. The asterisk (*) indicates that the final oxidase is unknown in ayahuasca.

Based upon our biosynthetic proposal, we hypothesized that the only missing component in HA biosynthesis was the generation of the tetrahydro-β-carboline scaffold. This scaffold formation could be enzyme-catalyzed or spontaneous, as tryptamine-derived molecules can undergo spontaneous Pictet-Spengler condensations with ketone or aldehyde substrates to yield tetrahydro-β-carboline scaffolds^27–31^. Critically, the installation of a hydroxy group at the C6 position on tryptamine (**1**), catalyzed by *Ph*T6H, activates this substrate for the Pictet-Spengler reaction^32,33^. Indeed, transient co-expression of *Ph*T6H with *Ph*TDC-1a in *N. benthamiana* yielded not only 6ʹ-hydroxytryptamine (**2**), but also low levels of 1-carboxy-tetrahydroharmol (**7**) and harmalol (**4**) (**Fig. 2g, SI Fig. 8**). The production of 1-carboxy-tetrahydroharmol (**7**) could result from the Pictet-Spengler condensation of 6ʹ-hydroxytryptamine (**2**) with endogenous pyruvate from *N. benthamiana*, which we observed to also occur spontaneously *in vitro* under acidic conditions (**Fig. 2g**, **SI Fig. 8**). Similarly, harmalol (**4**) production could occur via non-enzymatic oxidative decarboxylation of 1-carboxy-tetrahydroharmol (**7**), which also occurred spontaneously *in vitro* (**Fig. 2g**, **SI Fig. 8**). The production of 1-carboxy-tetrahydroharmol (**7**) and harmalol (**4**) provided a putative strategy to produce the HA scaffold in *N. benthamiana*. However, when we attempted to engineer the complete HA pathway using the five enzymes we had discovered (*Ph*TDC-1a, *Ph*T6H, *Ph*OMT-1, *Ph*FDO-1a, and *Ph*FDO-2), we produced only trace levels of harmaline (**5**) and no detectable harmine (**6**).

Our inability to reconstitute HA biosynthesis in *N. benthamiana* suggested that an enzyme may be required for HA scaffold formation. Therefore, we considered protein families previously established to catalyze this type of reaction, namely Bet v1/PR10 and strictosidine synthase families^34^, but did not identify any strong candidates based upon co-expression analysis. Therefore, we next explored non-enzymatic mechanisms that could facilitate formation of the HA scaffold, which can occur spontaneously through the condensation of 6ʹ-hydroxytryptamine (**2**) with either pyruvate (**SI Fig. 8**) or acetaldehyde (**SI Fig. 9**).

Radiolabeled ^14^C-pyruvate has been shown to be incorporated into the HA scaffold in *P. harmala*, presumably through a Pictet-Spengler condensation with 6ʹ-hydroxytryptamine (**2**) (**Fig. 2d**)^17,18^. However, this result is also consistent with the use of acetaldehyde as a co-substrate for scaffold formation, as decarboxylation of labeled pyruvate yields acetaldehyde with the same ^14^C-labeling pattern (**Fig. 2d**). Intriguingly, a putative pyruvate decarboxylase (*Ph*PDC-1), which would convert pyruvate into acetaldehyde^35^, was the 13^th^ highest co-expressing gene with *Ph*TDC-1a (**Fig. 2c, Supplementary Data 1**). Initial transient co-expression of *Ph*PDC-1 with *Ph*TDC-1a, *Ph*T6H, *Ph*OMT-1, *Ph*FDO-1a, and *Ph*FDO-2 in *N. benthamiana* did not impact HA biosynthesis (**SI Fig. 10**). However, sequence analysis revealed that *Ph*PDC-1 contains a plastid transit peptide (**SI Fig. 11**). This was unexpected, as canonical PDCs are generally cytosolic, where pyruvate pools are expected to be greater^36^. Indeed, sequence comparisons to other predicted PDCs from both *P. harmala* and public sequence databases did not identify another plant PDC with a predicted plastid transit peptide (**SI Fig. 11**). We hypothesized that localization of *Ph*PDC-1 was the limiting factor in our heterologous expression system, as no other HA biosynthetic enzymes were predicted to localize to the plastid (**Supplementary Data 2**). To test this, we co-expressed a known cytosolic PDC from the bacterium *Zymomonas mobilis* (*Zm*PDC)^37^ with all HA enzymes characterized thus far in *N. benthamiana*. This strategy resulted in harmine (**6**) biosynthesis (**SI Fig. 12**), supporting that the plastid localization of *Ph*PDC-1 may be the bottleneck in HA pathway reconstitution.

Using fluorescence microscopy, we found that fluorescently-tagged *Ph*PDC-1 localized to the plastid in *N. benthamiana* as predicted, and that truncating the signal peptide (*Ph*PDC-1Δ) re-localized this protein to the cytosol (**Fig. 2e**). Expression of *Ph*PDC-1Δ in *N. benthamiana* with *Ph*TDC-1a and *Ph*T6H led to the production of tetrahydroharmol (**3**) (**Fig. 2d**, **SI Fig. 9, 10**), and co-expression with the previously characterized HA biosynthetic enzymes led to the complete biosynthesis of harmine (**6**) (**Fig. 2d**, **SI Fig. 10**), supporting a role for *Ph*PDC-1 in generating acetaldehyde for scaffold formation. By including *Ph*PDC-1Δ, we observed reasonably high HA yields in *N. benthamiana* (185 µg harmaline (**5**) and 53 µg harmine (**6**) per gram dry leaf weight). Indeed, because harmaline (**5**) and harmine (**6**) each fluoresce when excited by ultraviolet (UV) radiation, we could directly visualize HA accumulation in *N. benthamiana* leaves under UV light (**Fig. 2f**).

The inability to reconstitute HA biosynthesis with the native *Ph*PDC-1 suggests a limitation in plastid pyruvate pools in *N. benthamiana* leaves, or a need for transport of HA intermediates in and out of the plastid. For HA biosynthetic reconstitution, either of these could be solved by re-localizing *Ph*PDC-1 to the cytosol. Regardless, we hypothesize that the noncanonical localization of *Ph*PDC-1 to plastids in *P. harmala* roots may serve to sequester reactive acetaldehyde while simultaneously providing a distinct compartment for HA scaffold formation to occur. Such a model would presumably require the import of both 6ʹ-hydroxytryptamine (**2**) and pyruvate into the plastid, and export of the tetrahydro-β-carboline product (**3**) back to the cytosol for the pathway to proceed. Although we were unable to identify functional transporters in this study, the unusual localization of *Ph*PDC-1 to plastids suggests a model of mixed compartmentalization for HA biosynthesis in *P. harmala*.

Encouraged by the high co-expression of the HA biosynthetic genes, we hypothesized that additional, unexpected biosynthetic components may be among the top correlating genes. Indeed, the third highest co-expressing gene to *Ph*TDC-1a shared homology with plant glyoxalase I (GLXI) enzymes (**Fig. 2c**). GLXI enzymes canonically act in the detoxification of methylglyoxal and other reactive aldehydes generated from cellular metabolism^38^. However, this GLXI-like enzyme from *P. harmala* (*Ph*GLXI-like) has modifications to active site residues that are highly conserved in canonical GLXI enzymes and instead shares greater homology with previously reported GLXI-like proteins from plants (**SI Fig. 13**)^39^. In support of a role in HA biosynthesis, transient co-expression of *Ph*GLXI-like with *Ph*TDC-1a and *Ph*T6H led to a 20-fold increase in harmalol (**4**) production (**Fig. 2g**). Furthermore, co-expression with the minimal set of biosynthetic genes (*Ph*TDC-1a, *Ph*T6H, *Ph*OMT-1, *Ph*FDO-1a, and *Ph*FDO-2) led to an additional 4-fold increase in harmaline (**5**) (**SI Fig. 14**). However, the effect from *Ph*GLXI-like was minimal when *Ph*PDC-1Δ was also included during pathway reconstruction, which could be explained by *Ph*PDC-1Δ having a much greater pathway boosting effect than *Ph*GLXI-like in the *N. benthamiana* heterologous system. Though a mechanism for the function of *Ph*GLXI-like in HA biosynthesis still needs to be elucidated, our results support a broader role for GLXI-like enzymes in plant metabolism, as previously hypothesized based on sequence analysis^39^.

Collectively, our work supports a model where HA scaffold formation in *P. harmala* can occur through two discrete, Pictet-Spengler reactions: 1) 6ʹ-hydroxytryptamine (**2**) condensation with acetaldehyde produced by *Ph*PDC-1, which we hypothesize to occur in plastids, and 2) 6ʹ-hydroxytryptamine (**2**) condensation with pyruvate in the cytosol, which seems to be influenced by *Ph*GLXI-like (**Fig. 2g**). We propose that both biosynthetic routes could contribute to HA production in *P. harmala* cells. Future work will allow us to determine the contribution of each pathway to HA biosynthesis and why *P. harmala* may utilize two routes for the same metabolic fate. We acknowledge that although the spontaneous condensation of 6ʹ-hydroxytryptamine (**2**) with either pyruvate or acetaldehyde occurs readily in *N. benthamiana* and *in vitro* (**SI Figs. 8, 9**), we cannot rule out that a dedicated enzyme catalyzes these scaffold forming reactions.

### Harmala alkaloid biosynthesis in *Banisteriopsis caapi*

Having uncovered the chemical logic of HA production in *P. harmala,* we next sought to elucidate HA biosynthesis in *B. caapi* and assess its evolutionary origins. We first profiled HA levels in tissues from mature *B. caapi* plants (**Fig. 3a**) and found that harmine (**6**) abundance was greatest in newly grown, aerial tissues (**Fig. 3b**). To profile gene expression across the *B. caapi* plant, we developed a tissue-specific RNA-seq dataset and used an in-house generated *B. caapi* draft genome for annotating protein-coding sequences. Using these data, we identified a single predicted TDC gene (*Bc*TDC) with enriched expression in the new, aerial tissues and confirmed its activity via transient expression in *N. benthamiana* (**SI Fig. 15**). This spatial pattern contrasted sharply with *P. harmala*, where active HA biosynthesis was concentrated in roots, thus providing an early indication that the two lineages deploy similar chemistry in distinct developmental and tissue contexts.

Using *Bc*TDC as a bait gene for co-expression analysis (**Supplementary Data 3**), we identified a CYP (*Bc*T6H; CYP71BG57) as the 4^th^-highest correlating gene (**Fig. 3c**). As in *P. harmala*, this CYP hydroxylates tryptamine (**1**) to form 6ʹ-hydroxytryptamine (**2**) (**Fig. 3d**). We next determined the function of multiple downstream tailoring enzymes whose expression correlated with *Bc*TDC in *B. caapi* (**Fig. 3c**). First, we identified two distinct *O*-methyltransferases (*Bc*OMT-1 and *Bc*OMT-2; 40% amino acid identity) that methylate HA intermediates. Specifically, *Bc*OMT-1 *O*-methylates tetrahydroharmol (**3**) to yield tetrahydroharmine (**8**) and harmalol (**4**) to yield harmaline (**5**), while *Bc*OMT-2 only acts on harmol (**9**) to produce harmine (**6**) (**Fig. 3d**, **SI Fig. 16**). Additionally, *Bc*OMT-1 can use 6ʹ-hydroxytryptamine (**2**) as a substrate to form 6ʹ-methoxytryptamine (**10**), a metabolite observed in *B. caapi* tissues (**SI Fig. 18**). Our co-expression analysis also identified an FDO from *B. caapi* (*Bc*FDO-1) that oxidizes tetrahydroharmol (**3**) to harmalol (**4**), and tetrahydroharmine (**8**) to harmaline (**5**) (**Fig. 3d**, **SI Fig. 17**). Despite testing numerous additional FDO and CYP candidates, we were unable to identify an enzyme that catalyzed the final oxidation to produce harmine (**6**). We hypothesize that this final oxidase either does not co-express strongly with the rest of the pathway genes or may belong to an unexpected enzyme family that was not prioritized here.

Informed by our findings on HA scaffold formation in *P. harmala*, we also searched the *B. caapi* transcriptome for genes encoding PDC or GLXI-like proteins. Genes with these annotations either did not co-express strongly with *Bc*TDC-1 or did not influence the HA pathway output when tested in *N. benthamiana*. However, the 7^th^ highest co-expressing transcript to *Bc*TDC-1 (out of 36,054 transcripts) was annotated as a threonine aldolase (TA), an enzyme that acts on *L*-threonine to produce acetaldehyde and glycine (**Fig. 3d**)^40^. Thus, we hypothesized that *B. caapi* may repurpose a threonine aldolase to supply acetaldehyde for HA scaffold formation. In support of this, transient co-expression of the putative threonine aldolase (*Bc*TA-1) with *Bc*TDC-1 and *Bc*T6H in *N. benthamiana* resulted in the production of tetrahydroharmol (**3**) (**Fig. 3d**). By combining *Bc*TA-1 with *Bc*TDC-1, *Bc*T6H, *Bc*FDO-1 and *Bc*OMT-1, we could reconstitute biosynthesis of harmalol (**4**), tetrahydroharmine (**8)**, and harmaline (**5**) in *N. benthamiana*, thus supporting a potential pathway for HA biosynthesis in *B. caapi* (**Fig. 3d**, **SI Fig. 18**).

These results suggest a plausible mechanism for HA scaffold formation in *B. caapi* through the non-enzymatic Pictet-Spengler condensation of threonine-derived acetaldehyde with 6ʹ-hydroxytryptamine (**2**) or 6ʹ-methoxytryptamine (**10**). Indeed, we found that carbons from isotopically labeled *L*-threonine were incorporated into harmaline (**5**) when this labeled substrate was fed to *B. caapi* plants (**Fig. 3e**). Although we presume the catalytic function of *Bc*TA-1 to be canonical, we found that *Bc*TA-1 was adjacent (60 kb) to another TA gene (*BcTA-2*) in the *B. caapi* genome, suggesting a duplication event (**Fig. 3f**). Indeed, phylogenetic analysis supports this duplication occurring more recently in *B. caapi* or a nearby related clade and not likely from a deeper ancestral duplication (**SI Fig 19**). In agreement with *Bc*TA-1 being specialized to HA biosynthesis, we found that *Bc*TA-2 expression did not correlate with *Bc*TDC-1 (rank 30,049) (**Fig. 3g**). We hypothesize that the ancestral TA gene in *B. caapi* was duplicated, and that one copy (*Bc*TA-1) gained novel regulation to co-express with HA biosynthetic genes. Thus, our results provide a plausible model by which *B. caapi* has adopted an independent strategy to supply acetaldehyde as a substrate for spontaneous Pictet-Spengler condensation with tryptamine-derived substrates in HA scaffold formation (**Fig 3d**). As with *P. harmala*, our results do not rule out the possibility that scaffold formation can be enzyme-catalyzed in *B. caapi*.

### Independent origins of harmala alkaloid biosynthesis in *P. harmala* and *B. caapi*

Having elucidated HA biosynthesis for both *P. harmala* and *B. caapi*, we next compared the evolutionary origins of the pathway enzymes in each species to ask to what extent each plant uses convergent or lineage-specific adaptations to produce HAs. The most compelling evidence for convergent evolution comes from the T6H genes, which arose independently in each lineage through duplication of an ancestral tryptamine 5ʹ-hydroxylase (T5H) gene (**Fig. 4a, b**). In the genome of *B. caapi*, the *Bc*T6H gene is 50 kb from its closest paralog, a functional T5H (hereafter, *Bc*T5H; CYP71BG57) (**Fig. 4b-d**). This suggests that *Bc*T6H may have arisen through the duplication and neofunctionalization of *Bc*T5H. With this result in mind, we revisited the evolutionary origins of the T6H gene from *P. harmala*. First, we identified and biochemically characterized a T5H (*Ph*T5H; CYP71BG48) from the *P. harmala* transcriptome, which shares high amino acid sequence identity (79%) with *Ph*T6H (**Fig. 4b, c**). Next, we found that the T6H ortholog in *P. multisectum* is also in genomic proximity (6.8 kb) to its paralogous T5H gene (**Fig. 4d**), suggesting a convergent evolutionary mechanism for the origin of T6H in both species.

Though relatively few T5H genes from plants have been biochemically characterized, their activity is implicated in the production of key metabolites, namely serotonin and melatonin^41–44^. These metabolites are found in many angiosperm lineages^45,46^, suggesting that T5H may be broadly conserved. Indeed, BLAST searches identified potential T5H orthologs from plant species throughout the angiosperms (**Fig. 4b**, **SI. Fig. 20**). To assess the conserved function of these genes, we identified and functionally characterized T5H orthologs from eight phylogenetically dispersed plant orders (Asterales, Caryophyllales, Fagales, Malpighiales, Malvales, Ranunculales, Rosales, Sapindales), thus supporting that T5H genes are widespread in angiosperms, though not universal (**Fig. 4b**, **SI. Fig. 20-21**). Together, these data suggest a model in which analogous T6H genes arose in *P. harmala* and *B. caapi* through the independent duplication of an ancestral T5H from plant serotonin metabolism, as demonstrated by phylogenetic analyses that show *Bc*T6H and *Ph*T6H are each more closely related to their T5H paralog than to each other (**Fig. 4b, SI Fig. 20**).

The switch from T5H to T6H activity requires a minor shift in the location of the installed hydroxy group on tryptamine, suggesting that only a few active site modifications could alter function. Therefore, we next considered which amino acid changes conferred a change from C5 to C6 hydroxylation regiospecificity. Using structural modeling and sequence alignments, we identified four residues in proximity to the active site of both T6Hs that differed from the conserved T5H sequence (**SI Fig. 22-23**). Notably, the specific changes identified near the active sites of *Ph*T6H and *Bc*T6H are distinct (**SI Fig. 23**), supporting independent neofunctionalization events for these enzymes within the context of a broader evolutionary convergence. We next performed site directed mutagenesis with each T6H to substitute the four selected amino acids with the conserved T5H residues. Although a single mutation in *Ph*T6H (T490A) could confer some T5H activity, the analogous change in *Bc*T6H (S483A) led to total loss in hydroxylase function. However, modifying all four selected amino acids in *Ph*T6H and *Bc*T6H to the ancestral T5H residues resulted in a complete shift to T5H activity in both enzymes (**SI Fig. 24**). Thus, our results indicate that while T6H function may have arisen from an ancestral T5H gene in both *P. harmala* and *B. caapi*, a distinct set of amino acid substitutions led to this functional convergence.

We also evaluated the evolutionary origins of the downstream HA methyltransferase and oxidase enzymes from *P. harmala* and *B. caapi* that have converged on analogous biosynthetic chemistry. First, we note that *O*-methyltransferase (OMT) function differs between *B. caapi* and *P. harmala*. While *Bc*OMT-1 and *Bc*OMT-2 exhibit limited substrate specificity, a single, promiscuous methyltransferase in *P. harmala* (*Ph*OMT-1) can catalyze the *O-*methylation of three HA substrates (**SI Fig. 5, 16**). *Bc*OMT-1 and *Bc*OMT-2 share low amino acid sequence identity with *Ph*OMT-1 (43% and 40%, respectively) and phylogenetic analysis demonstrated that these *O-*methyltransferases cluster into distinct clades (**SI Fig. 25**), supporting independent origins of OMT function between *P. harmala* and *B. caapi*. Similar evidence for independent evolution was observed for the downstream oxidase enzymes. Specifically, *Bc*FDO-1 shares less than 60% amino acid identity with the *Ph*FDO-1 homologs, and phylogenetic analysis indicates that these analogous oxidase genes also arose independently in a lineage-specific manner (**SI Fig. 26**). Collectively, although the scaffold tailoring reactions are catalyzed by the same enzyme families in *P. harmala* and *B. caapi*, our results show that the individual enzymes evolved in parallel in these plants.

Overall, the chemical logic of HA biosynthesis is nearly identical in *P. harmala* and *B. caapi* (**SI Fig. 27**), with pathway enzymes in the two species arising independently through duplications and neofunctionalization of similar ancestral protein families, or in the case of T6H, from functionally identical T5H enzymes (**Fig. 4e**). Except for the TDCs, which seem to be widespread and ancestral in angiosperms^23^, our results support a model where HA biosynthetic genes in *P. harmala* and *B. caapi* arose in parallel, but primarily through convergent evolutionary and functional mechanisms involving neofunctionalization of a limited set of ancestral enzymes. In both species, the same enzyme families were recruited for C6 hydroxylation, *O*-methylation, and downstream oxidations. In contrast, how acetaldehyde is putatively generated for scaffold formation appears to be lineage specific. In theory, acetaldehyde could be generated through several primary metabolic routes that are conserved in plants^35,40,47,48^. Our data suggest that *P. harmala* and *B. caapi* have co-opted distinct pathways from primary metabolism to produce acetaldehyde. These proposed modes of acetaldehyde biosynthesis do not require novel enzyme catalysis, but rather non-canonical cellular localization (*Ph*PDC-1) or gene duplication and modified expression of a paralog (*Bc*TA-1). Collectively, our work uncovers a striking pattern of biosynthetic convergence in *P. harmala* and *B. caapi*, with both parallel recruitment of the same enzyme families and co-option of distinct enzymes from primary metabolism to achieve HA production.

## Discussion

We have elucidated the molecular basis for HA biosynthesis in *P. harmala* and *B. caapi*, two neuroactive plants that have had a critical role in medicine and ritual across continents, climates, and cultures for millennia. Though several routes to HAs are possible, we find that these plants have converged on HA production through remarkably similar genetic and biochemical processes, including independently evolved strategies to produce 6ʹ-hydroxytryptamine (**2**) and acetaldehyde, the key building blocks of the HA scaffold. The most notable similarity between the two HA pathways is the independent duplication and neofunctionalization of an ancestral T5H from serotonin metabolism to generate T6H, the enzyme that produces 6ʹ-hydroxytryptamine (**2**). Although the change in function between T5H and T6H is subtle, the location of the hydroxy group in 6ʹ-hydroxytryptamine (**2**) makes this substrate more reactive than serotonin for condensation with acetaldehyde^32,33^. Thus, T6H catalyzes a committed step in HA biosynthesis by producing a substrate that is primed for a rapid, non-enzymatic Pictet-Spengler reaction. This scaffold-generating condensation also requires an increased concentration of acetaldehyde in plant cells, which *P. harmala* and *B. caapi* appear to have achieved through the independent recruitment of distinct enzymes from primary metabolism. Together, our results reveal a repeating theme in plant HA biosynthesis, where modest changes to pre-existing biosynthetic machinery can yield the reactive intermediates needed to form the HA scaffold.

In conclusion, *P. harmala* and *B. caapi* illustrate convergence across an exceptionally broad hierarchy. Independently adopted for medicine and ritual by distinct cultures spanning from Eurasia and North Africa to Amazonia, these distantly related plants also converge on identical neuroactive metabolites. Although these plants use remarkably similar biochemistry to synthesize HAs, we reveal that the underlying enzymes arose through a combination of parallel evolution and lineage-specific innovation. The recurrence of similar biosynthetic solutions, including the neofunctionalization of serotonin metabolic enzymes and use of non-enzymatic scaffold-forming chemistry, suggests that the pre-existing metabolic repertoires of both plants were conducive to the evolution of HA biosynthesis. Our results demonstrate that each plant independently recruited these similar ancestral metabolic processes in parallel to achieve the same biochemical outcome: the production of neuroactive HAs. Collectively, our findings show how comparative studies of medicinal plants with identical bioactive molecules can integrate ethnopharmacology with biochemistry and genomics to reveal how plant chemistry evolves and may be constrained, and how humans have repeatedly recognized and incorporated its biological effects.

## METHODS

### Chemicals and reagents

All chemicals and reagents were obtained from commercial vendors, unless otherwise noted. Authentic standards for harmalol (**4**) (hydrochloride hydrate), tetrahydroharmine (**8**), harmaline (**5**), and harmine (**6**) were obtained from Cayman Chemicals. Harmol (**9**) was purchased from Toronto Research Chemicals and 6ʹ-hydroxytryptamine (**2**) from Enamine. Sodium pyruvate and acetaldehyde were obtained from Millipore Sigma. The following labeled substrates were obtained from Cambridge Isotope Laboratories: *L*-tryptophan (indole-D₅), *L*-threonine (4-¹³C, 2,3-D₂), sodium borodeuteride-D_4_, and sodium pyruvate (2,3-¹³C₂).

### D_2_-tetrahydroharmol (3) synthesis

D_2_-tetrahydroharmol (**3**) was synthesized by the imine reduction of harmalol (**4**) using sodium borodeuteride-D_4_ (NaBD_4_) as a reducing agent based on a modified protocol^49^. Specifically, 0.1 mmol of harmalol (**4**) hydrochloride hydrate was dissolved in 1 mL of cold methanol (HPLC-grade) and mixed with 0.1 mmol of NaBD_4_ previously dissolved in 100 µL of cold methanol (HPLC-grade). The reaction mixture was stirred for 30 min at 4 ℃ and subsequently dried under a nitrogen gas stream.

### Tetrahydroharmol (3) synthesis

Tetrahydroharmol (**3**) was synthesized via the Pictet-Spengler reaction of 6ʹ-hydroxytryptamine with acetaldehyde using a modified protocol^50^. Due to the volatility of acetaldehyde, an excess of this reagent was used to ensure the reaction proceeded forward. A 200 µL reaction was prepared by mixing 175 µL of 125 mM *L*-tartaric acid, 20 µL of 50 mM 6ʹ-hydroxytryptamine (**2**) (in DMSO), and 5 µL 1 M acetaldehyde and incubated in a water bath at 60 ℃ for 1.5 h. Reaction products were diluted in 80% methanol (1:200) and analyzed by LC-MS.

### 1-carboxy-tetrahydroharmol (7) and ¹³C₂-carboxy-tetrahydroharmol synthesis

The Pictet-Spengler reaction of 6ʹ-hydroxytryptamine (**2**) and pyruvate was used to generate 7-hydroxy-1-methyl-1,2,3,4-tetrahydro-β-carboline-1-carboxylic acid (1-carboxy-tetrahydroharmol (**7**)). To aid with structural validation the same reaction was prepared with sodium pyruvate (2,3-¹³C₂) to produce ¹³C₂-carboxy-tetrahydroharmol (**7**). A modified protocol^50^ was used for both reactions. A 200 µL reaction was prepared by mixing 170 µL of 125 mM *L*-tartaric acid, 20 µL of 50 mM 6ʹ-hydroxytryptamine (**2**) (in DMSO), and 10 µL of either 100 mM sodium pyruvate or its labeled form (2,3-¹³C₂). Reactions were incubated at room temperature for 2 h and diluted in 80% methanol (1:20) for LC-MS analysis.

### Plant growth

*Peganum harmala* seeds (Companion plants) were surface sterilized and sown in 0.5 X MES, pH 5.8, 0.7% (w/v) agar and stratified for two days. Seedlings germinated at room temperature under long-day conditions (16 h light/ 8 h dark). Upon germination (15 days) plants were transferred to 0.5 X Murashige and Skoog (MS) medium, pH 5.8, 1% (w/v) sucrose, 0.7% (w/v) agar and grown vertically. *Banisteriopsis caapi* plants were grown in a glasshouse in the Biological Laboratories building and in a grow room in Weld Hill Growth Facilities (Arnold Arboretum). Cuttings of *B. caapi* were periodically transplanted into new soil to initiate new shoot growth.

*Nicotiana benthamiana* seeds were sown in Pro-Mix HP AGTIV Reach soil soaked with a fertilizer solution (Southern Ag PowerPak 20-20-20 Water Soluble Fertilizer with micronutrients) and were germinated in growth chambers at 30 ℃ with a 16 h light/ 8 h dark cycle. Seedlings were kept under humidity domes for one week, after which domes were removed for the remaining growth. Plants were used for Agrobacterium mediated transient expression experiments 4-5 weeks after germination. Following infiltration/transformation, plants were grown at room temperature with the same light/dark cycle.

### Isotope tracer studies in *P. harmala* and *B. caapi*

To track active biosynthesis in *P. harmala*, we initially attempted to feed mature plants with D_5_-tryptophan, but the label was not incorporated evenly throughout the plant. Therefore, we opted to use seedlings, which we suspected would be more amenable to uptake of labeled tryptophan due to their smaller size and active growth. Twenty-four-days-old *P. harmala* seedlings were transferred to MS liquid medium supplemented with either DMSO, 1 mM *L*-tryptophan, or 1 mM D_5_-tryptophan for 48 h. Seedlings were rinsed, tapped dry, flash-frozen, and homogenized under liquid nitrogen. Metabolites were extracted in 80% methanol (20 µL mg^-1^ fresh weight) for 2 h at room temperature. Extracts were filtered through 0.45 µm PTFE membrane filters and diluted in 80% methanol prior to analysis via LC-MS.

For isotope tracer experiments in *B. caapi* plants, new growth tissues including a ∼5 cm stem and two new leaves were excised and placed in vials containing 1700 µL of water, 1 mM D_5_-tryptophan, or 1 mM *L*-threonine (4-¹³C, 2,3-D₂). Plants were kept in the greenhouse covered with domes for 74 h. During this time the feeding solution was refilled as needed. After this time, samples were tapped dry, flash-frozen in liquid N_2_, lyophilized to dryness, and homogenized with a ball mill. Metabolites were extracted in 80% methanol (80 µL mg^-1^ fresh weight) and prepared for LC-MS analysis as described above. Unlike labeled *L*-threonine, we were unable to detect D_5_-tryptophan uptake or incorporation into HAs in *B. caapi* cuttings.

### LC-MS analysis

All samples were analyzed on an Agilent 1290 Infinity II ultrahigh-performance liquid chromatography (UHPLC) system coupled to a 6546 quadrupole time of flight (QTOF) mass spectrometer (Agilent Technologies, Santa Clara, CA) with the specific parameters indicated in **Supplementary Methods**. In general, analytes were separated using an Agilent ZORBAX RRHD Eclipse Plus C18 column (2.1 x 50 mm, 1.8 µm) using a gradient method with water with 0.1% (v/v) formic acid and acetonitrile with 0.1% (v/v) formic acid as the mobile phase buffers. Electrospray ionization (ESI) in positive mode was used for detecting all metabolites in this study.

Tandem mass fragmentation (MS/MS) with collision energies of 10, 20, 40 V were applied to plant extracts and standards to obtain structural information and verify the identity of metabolites. Data analysis was performed with both the Agilent Mass Hunter Qualitative (version 10.0) and Quantitative (version 11.0) software. Untargeted data analysis was performed with XCMS (Scripps Center for Metabolomics) with data converted to mzML format using MSConvert (ProteoWizard) as an input with default parameters for positive ionization mode. A previously described R script^21^ was utilized for pairwise comparisons. All graphs and statistical analysis were performed in R studio (R version 4.2.0).

### *Banisteriopsis caapi* plant material, DNA extraction, genome sequencing, and assembly

Living *Banisteriopsis caapi* material was collected from plants maintained at The Kampong National Tropical Botanical Garden in Coconut Grove, Florida, USA. Fresh young stem and leaf tissues were collected, flash frozen using liquid nitrogen, and stored at −80 °C prior to DNA extraction. High molecular weight genomic DNA was extracted following the general approach used previously for plant genome sequencing in the Davis laboratory^51^. Briefly, frozen tissue was ground under liquid nitrogen, nuclei-enriched material was isolated to reduce organellar DNA and secondary metabolites, and genomic DNA was purified using a CTAB-based extraction. The resulting high molecular weight DNA was cleaned using 3X KAPA Pure beads (Roche Sequencing Solutions) and assessed for concentration and quality prior to library preparation. DNA purity was assessed using a Nanodrop spectrophotometer (Thermo Fisher Scientific) and concentration was measured with a Qubit fluorometer (Thermo Fisher Scientific). DNA size was assessed using a Tapestation system (Agilent).

For genome sequencing, we applied a combination of long and short read sequencing. High molecular weight DNA from *B. caapi* was used to generate PacBio HiFi long-read data using a PacBio Sequel instrument following standard manufacturer protocols. We also generated Illumina short-read genomic data sequenced on Illumina Novaseq S4, 2×150 flowcell, which were used for k-mer–based assembly evaluation.

The *B. caapi* genome was assembled using a Snakemake-based PacBio HiFi assembly workflow (https://github.com/harvardinformatics/pacbio_hifi_assembly), which implements hifiasm v0.19.9 for *de novo* assembly^52^. Assembly statistics were calculated using QUAST^53^. The resulting draft assembly was 1.2 Gb in total length and consisted of 558 contigs, with a contig N50 of 9.9 Mb. Genome completeness and consensus accuracy were assessed with Merqury v1.3 using k-mers derived from the Illumina short-read data (k = 20)^54^. This analysis estimated the assembly to be 89% complete, with an average consensus quality score of Q61. We also estimated genome completeness using the compleasm software package (v0.2.8) to annotate BUSCO genes using the eudicotyledons_odb12 dataset, which gave an estimated completeness of 99.6% (88.5% single copy, 10.8% duplicate).

Repetitive elements were identified de novo using the Dfam TETools toolkit v1.89.2 (https://github.com/Dfam-consortium)^55^. RepeatModeler was used to generate a *B. caapi*–specific repeat library^56^, and RepeatMasker was then used to annotate repetitive sequences in the genome assembly^57^. Approximately 75% of the assembled genome was annotated as repetitive sequence. This consists almost exclusively of interspersed elements, with retroelements making up 46.4% of repeats (specifically LTR elements with 41.9%), 4.4% DNA transposons and 23.5% other unclassified interspersed elements.

We also obtained tissue-specific RNA-seq data to guide gene prediction and annotation. For this, total RNA for each tissue was extracted from single replicates of four *B. caapi* tissue types (leaf, stem, root, apical meristem) using the Monarch Total RNA Miniprep kit (New England Biolabs). The concentration, quality, and integrity of the RNA samples were examined using Qubit, Nanodrop, and Tapestation. cDNA libraries were prepared using KAPA mRNA HyperPrep kit (Roche Sequencing Solutions). The samples were then sequenced on Illumina NovaSeq S4, 2×150 lane. These RNA-seq data were used to construct a reference transcriptome, which was generated using a previously outlined workflow^58^ that utilizes StringTie v. 2.1.2 for transcript assembly^59^. A separate RNA-seq data set was also subsequently collected for the explicit purpose of gene co-expression analysis (described in detail below). The resulting draft genome was used together with the two tissue-specific RNA-seq data sets to support discovery and validation of candidate HA biosynthetic genes presented in the main text, including assessment of local genomic context for duplicated loci. Because genome-wide structural gene annotation was not performed for this assembly, candidate coding sequences were primarily inferred from transcriptomic data and annotated using protein-domain and similarity searches as described below.

### RNA-seq for gene expression profiling of *P. harmala* and *B. caapi* tissues

Based upon isotope tracer experiments, four twenty-six-days-old *P. harmala* seedlings were harvested and dissected into cotyledons, leaves, stems, and roots (**Fig. 2a**). Roots were further sectioned into three parts of equal length: root top (basal region), root middle, and root tip (apical region). Tissues were flash frozen and stored at -80 °C until the day of RNA extraction. Tissue was homogenized by mortar and pestle and RNA was extracted with the QIAGEN RNeasy® Plant Kit and treated with DNAse using the Invitrogen™ TURBO DNA-free™ kit following manufacturer’s instructions. RNA quantification and quality control were performed by Azenta Life Sciences using RNA Qubit and TapeStation. Library preparation was performed by Azenta Life Sciences with NEBNext Ultra II RNA Library Preparation Kit with Poly-A Selection (New England Biolabs) and sequenced on an Illumina NovaSeq X Plus, 2×150bp lane.

Using metabolite distribution as a guide, four one-year-old *B. caapi* plants were harvested and dissected into lateral roots, main roots, lateral stems, main stems, young leaves, petioles, new leaves and shoot apical meristem (**Fig 3a**). RNA extraction, library preparation, and RNA sequencing were performed following the same methods as those described for *P. harmala*.

We generated a reference transcriptome of full-length transcripts for *P. harmala* using PacBio Isoseq™, with cDNA library preparation performed by the Bauer Core Facility at Harvard University. Equimolar RNA quantities from the 24 samples described above were mixed and used for library preparation using the NEBNext Single Cell/Low Input cDNA Synthesis and Amplification Module kit (New England Biolabs) and the Iso-Seq v2 SMRTbell prep kit 3.0 (PacBio) following manufacturer’s instructions. The library was sequenced on a SMRT Cell 8M on a Sequel IIe system (PacBio). Isoseq reads were processed with the isoseq package v4.0.0 (PacBio). First, the cDNA adapters added during library preparation were removed with the command lima, then polyA tails and artificial concatemers were removed with the isoseq refine command from the same package. Reads were clustered based on the presence of at least two reads per transcript with the isoseq cluster2 command, which resulted in 223,253 reads. Reads were further collapsed using a 99% sequence similarity threshold using the package CD-HIT v4.8.1^60^ (parameters: cd-hit-est -I <> -o <> -c 0.99 -n 11 -d 500 -M 10000 -T 10) which resulted in 73,698 transcripts.

Illumina reads from both species were processed with trimmomatic v0.39^61^, hard-clipped for the first 14 base pairs, trimmed for adapters and low-quality reads (parameters: ILLUMINACLIP:TruSeq3-PE.fa:2:30:10 HEADCROP:14 LEADING:5 SLIDINGWINDOW:4:5 MINLEN:50). For *P. harmala,* transcript abundance was quantified based on a pseudoalignment between the collapsed Isoseq transcriptome and the trimmed Illumina reads using the package kallisto v0.48.0 with default parameters^62^, generating a counts matrix. Normalized expression values as Trimmed Mean of M (TMM)-normalized transcripts per million were estimated using a script (abundance_estimates_to_matrix.pl) from the Trinity v2.15.2 package^63^. Transcript abundance for *B. caapi* was determined using the same pseudoalignment methods described above using the *B. caapi* trimmed Illumina reads and the genome-guided *B. caapi* transcriptome as a reference. Sample clustering analysis of the *B. caapi* samples revealed that unlike all other samples, the shoot apical meristem samples did not form a cluster, therefore they were excluded from downstream analyses, including co-expression analysis.

For both *P. harmala* and *B. caapi* transcriptomes, coding regions for each transcript were identified using TransDecoder v5.7.0 (https://github.com/TransDecoder/Transdecoder/). To annotate the transcriptome, the peptide sequences from the coding sequences predicted above were queried against the protein family database PFAM using hmmscan with default parameters from the HMMER v3.3.2 package^64^. To complement the annotation, the peptide sequences were compared to the UniProtKB/Swiss-Prot reviewed, non-redundant, protein sequence database (niport.org) using the blastx command from the blast-plus v2.13.0 package^65^.

The Pearson’s correlation co-efficient for all genes relative to either *Ph*TDC-1a and *Ph*TDC-1b or *Bc*TDC were calculated using the TMM-normalized reads and the cor() function in rStudio (R version 4.2.0). Gene candidates were selected based on high correlation rankings to bait genes as well as their biosynthetic relevance based on PFAM and BLAST hit annotations.

### Candidate biosynthetic gene cloning

For cDNA synthesis, the same RNA samples used for RNA-seq proceeding from tissues with the highest TDC expression were selected (a root mid sample for *P. harmala* and a new leaf for *B. caapi*) and reverse transcribed using the SuperScript™ III First-Strand Synthesis SuperMix kit (Invitrogen) with oligo(dT)_20_ primers following manufacturer’s instructions. Coding regions for candidate genes were either synthesized or PCR amplified with Q5® High Fidelity DNA Polymerase (New England Biolabs) from cDNA using gene-specific primers with 5ʹ and 3ʹ overhangs compatible with the pEAQ-HT vector^24^. PCR products were run on an agarose gel, excised, and purified with the Zymoclean Gel DNA Recovery Kit (Zymo Research). The pEAQ-HT vector was linearized with restriction enzymes *Age*I and *Xho*I, and candidate genes were cloned into the vector with NEBuilder® HiFi DNA Assembly Master Mix (New England Biolabs), a Gibson assembly-based system. *E. coli* 10-beta competent cells (New England Biolabs) were transformed with the assemblies and plated on Luria-Bertani (LB) media with kanamycin selection (50 µg mL^-1^). The presence of an insert in the *E. coli* colonies was verified by colony PCR using plasmid-specific primers. Plasmids were extracted from liquid cultures with the ZR Plasmid Miniprep (Zymo Research) and sent for either Sanger (Azenta Life Sciences) or whole-plasmid sequencing (Plasmidsaurus). Mutated versions of *Ph*T5H, *Ph*T6H, *Bc*T5H, and *Bc*T6H were synthesized with appropriate Gibson overhangs (Twist Biosciences) and cloned into the pEAQ-HT plasmid as described above.

### Signal peptide prediction and cloning of truncated *Ph*PDC-1

*In silico* protein localization for *P. harmala* protein sequences was predicted using TargetP 2.0^66^ (https://services.healthtech.dtu.dk/services/TargetP-2.0/) and reported in **Supplementary Data 2**. The *Ph*PDC-1 protein sequence was analyzed using both TargetP 2.0 and LOCALIZER^67^ (https://localizer.csiro.au) for the presence of an N-terminal signal peptide. Both algorithms predicted a plastid transit peptide, although of different lengths. Accordingly, we generated two truncated versions of *Ph*PDC-1, either missing 41 or 47 amino acids at the N-terminus and after functional testing by transient expression in *N. benthamiana* we found that both were functional. *Ph*PDC-1(-47a.a.) had a greater effect than *Ph*PDC-1(-41a.a.) on overall pathway output, therefore we opted to move forward with *Ph*PDC-1(-47 a.a.) for further analyses, hereafter referred as *Ph*PDC-1Δ.

### Cloning of fluorescently-tagged proteins

Gene fragment fusions with *Ph*PDC-1 or *Ph*PDC-1Δ and C-terminus mNeonGreen^68^ were synthesized by Twist Biosciences. The fragments consisted of either *Ph*PDC-1 or *Ph*PDC-1Δ without a stop codon, a 12bp linker (GGTTCGGGAAGC), and mNeonGreen without a start codon. The gene encoding for the fluorescent protein was codon optimized for expression in *N. benthamiana*. Gene fragments contained 5ʹ and 3ʹ overhangs compatible with the pEAQ-NOS vector. Gene fusions were cloned into pEAQ-NOS, a vector generated by the Sattely Lab (Stanford University) by substituting the CaMV35S promoter with the NOS promoter. The pEAQ-NOS vector was selected for localization experiments to prevent localization artifacts caused by the high overexpression resulting from the CaMV35S promoter in pEAQ-HT^24^. Cloning was performed as described under “Candidate biosynthetic gene cloning.”

### Transient gene expression in *Nicotiana benthamiana* leaves and metabolite extraction

*Agrobacterium tumefaciens* strain GV3101 was transformed with a pEAQ-HT^24^ plasmid carrying the gene of interest and grown on LB with kanamycin (50 µg mL^-1^) and gentamycin (30 µg mL^-1^) selection. The presence of an insert in the Agrobacterium colonies was verified by colony PCR using plasmid-specific primers. Liquid cultures were grown for 24 h, after which cells were harvested by centrifugation, rinsed with LB, and induced with 10 mM MES buffer pH 5.6, 10 mM MgCl_2_, 150 µM acetosyringone for 2 h. The OD_600_ was measured for each culture and diluted to OD_600_ = 0.3 with induction media. Four-weeks-old *Nicotiana benthamiana* leaves were infiltrated with the cultures and incubated at room temperature under long-day conditions (16 h light/ 8 h dark) for 3-5 days. For substrate infiltration, substrates diluted to 50-100 µM with water were infiltrated into the infected zone and incubated for 24 h. The infection zone from each leaf was excised, flash frozen, and lyophilized overnight. Samples were homogenized with a ball mill and metabolites were extracted and analyzed as described above.

### Fluorescent protein localization via confocal microscopy

Using the Agrobacterium-mediated transient expression system, *N. benthamiana* leaves expressing C-terminal mNeonGreen fusion proteins were harvested 96 h post-infiltration. Approximately 1 cm x 1 cm leaf tissues were excised from the infiltrated area and mounted in water abaxial-side up under 22 mm cover slips and sealed inside with melted beeswax. These were then imaged by confocal microscopy using autofluorescence in chloroplasts as a co-localization marker.

Fluorescence images were acquired on a Zeiss LSM 880 confocal microscope mounted on an AxioObserver stand using a LD C-Apochromat 40x/1.1 W Korr M27 water-immersion objective and GaAsP photomultiplier tube detectors. mNeonGreen was excited at 488 nm with a detection window of 499-536 nm. Chlorophyll autofluorescence was excited simultaneously at 488 nm with a detection window of 663-735 nm. Transmitted-light images were acquired during confocal scanning with the 488 nm laser serving as the illumination using a transmitted-light photomultiplier detector (T-PMT). The pinhole was set to 1.0 Airy unit for all channels. Images were acquired with a pixel dwell time of 0.552 μs, 2x line averaging, and a z-step size of 1.0 μm. The Z-stack range was selected to encompass the cell of interest resulting in varied numbers of optical sections among images.

ImageJ (version 1.54p, Fiji 2.16.0^69^) software (https://imagej.net/) was used to process Z-stacks and generate maximum intensity projections. For displaying images, brightness and contrast were adjusted in ImageJ using auto settings within the B&C window.

### Phylogenetic analysis

The protein sequences used to populate the phylogenetic trees were identified by BLASTP (https://blast.ncbi.nlm.nih.gov) searches against the Viridiplantae ClusteredNR database using *P. harmala* and *B. caapi* protein sequences as queries. The top 100 hits for each sequence were selected and redundant hits among independent BLASTp searches were removed using the package CD-HIT v4.8.1 (parameters: cd-hit -c 1.0 -n 5)^60^. Multiple sequence alignments were computed using the package MAFFT v7.525 (parameters: mafft --localpair --maxiterate 3000 -- thread 32)^70^. The alignment generated for SI Fig. 22 was visualized using ESPript 3.2 (https://espript.ibcp.fr)^71^. Phylogenetic trees were generated using the protein alignments as inputs for IQ-TREE multicore v2.3.6 (parameters: iqtree -m MFP -keep_empty_seq --no-terrace -T AUTO -bb 1000 -alrt 1000 -bnni)^72–74^. For Fig. 4b the alternative tree topology where *Bc*T5H clades with *Ph*T5H and *Bc*T6H clades with *Ph*T6H was evaluated using the approximately unbiased (AU) test^75^ performed with 10,000 resamplings using the RELL method available in the IQ-TREE package. The alternative tree topology was rejected (*p* = 2.79e-45). Trees were visualized and color-coded based on plant order using iTOL v7 (https://itol.embl.de)^76^.

### Transcript, protein, and genome sequence analyses

Routine visualization and comparison of candidate gene transcript and protein sequences was performed using Geneious Pro software. This includes prediction of signaling peptides, prediction of transmembrane domains, and multiple sequence alignments, which could all be performed using available plugins. Predicted protein structures for *Ph*T5H, *Ph*T6H, *Bc*T5H, and *Bc*T6H were generated using the AlphaFold server (https://alphafoldserver.com/). Each structure was generated with heme as a bound ligand. Protein structures were visualized using PyMOL.

Analysis of the *P. multisectum*^25^ and *B. caapi* genomes was performed using Geneious Pro software. The locations of biosynthetic pathway genes on these genomes were identified by BLASTN searches and by comparing relative chromosomal locations from General Feature Format (GFF) files. Dot plots comparing various loci were generated through multiple sequence alignments of a locus against itself. Sequence similarity was subsequently visualized using a word size of 20.

### Images and geographic distribution of *B. caapi* and *P. harmala*

The image of *P. harmala* in **Fig. 1a** was taken by Andreas Manz and is available under a CC0 license on iNaturalist (https://www.inaturalist.org/observations/278909553). The image of *B. caapi* in **Fig. 1a** is available under a CC0 license from Wikimedia Commons (https://commons.wikimedia.org/wiki/File%3ABanisteriopsis_caapi_flower.jpg) and was provided by the author Maria Gabriela Yejo. Other plant images (**Figs. 2 & 3**) were taken by the authors. The plot of *B. caapi* and *P. harmala* geographic distributions (**Fig. 1b**) was generated by compiling the documented observations of these plants from the Global Biodiversity Information Facility (GBIF; gbif.org). We then used Plants of the World Online (powo.science.kew.org) as a cross-reference so that we only included observations from countries to which each plant is endemic. The geographic coordinates (latitude/longitude) of each observation were then plotted on a Mollweide projection of the world using the sf, rnaturalearth, and rnaturalearth data packages in R.

### Data analysis and statistical information

For transient expression experiments in *N. benthamiana*, at least three biological replicates were routinely used for gene characterization. Replicates consisted of independently-transformed leaves from different plants. All statistical tests (*t*-tests and one-way ANOVAs) were two-tailed and performed in R.

## Data availability

The materials and data generated from the work in this manuscript will be made available upon request. The raw RNA-seq reads from *P. harmala* and raw RNA-seq and genomic reads for *B. caapi* will be deposited to the NCBI Sequence Read Archive (SRA). The assembled transcriptomes (*P. harmala* and *B. caapi*) and assembled genome contigs (*B. caapi*) will be deposited to GenBank. Individual nucleotide and protein sequences for the genes characterized in this study will also be deposited to GenBank. All sequences will be made available following publication.

## Acknowledgements

We thank the staff from the Biological Laboratories Plant Growth Facilities and Weld Hill Growth Facilities (Harvard University) for helping with the growth of *P. harmala* and *B. caapi*. Fairchild Botanical Garden and The Kampong National Tropical Botanical Garden supplied fresh materials for *B. caapi*. Thank you to the Harvard Center for Biological Imaging (HCBI) for assistance with fluorescence microscopy. The computations in this paper were run on the FASRC Cannon cluster supported by the FAS Division of Science Research Computing Group at Harvard University. We acknowledge Prof. George Lomonossoff (John Innes Centre) for providing the pEAQ-HT plasmid and Prof. Elizabeth Sattely (Stanford University) for providing the pEAQ-Nos plasmid. We thank Prof. David Nelson (University of Tennessee) for aiding with cytochrome P450 nomenclature. We’re also grateful to Andrea Loya Castillo and Liam Joy for their assistance with gene cloning.

## Author Contributions

The project was conceptualized by FMV, CCD, and RSN. All experimental work related to the discovery of HA biosynthesis in *P. harmala* and *B. caapi* was carried out by FMV, with assistance from YWH and JAO. Experiments, data analysis, and writing relevant to the transcriptome and genome sequencing of *B. caapi* were performed by JW, YY, DEK, TBS, and CCD. FMV and RSN wrote the manuscript with contributions from TBS and CCD.

## Other data

- Supplementary Information
- Supplementary Data 1 - Unfiltered table of *Peganum harmala* transcripts co-expressed with *Ph*TDC-1a.
- Supplementary Data 2 - *In silico* predicted localization of *Peganum harmala* enzymes.
- Supplementary Data 3 - Unfiltered table of *Banisteriopsis caapi* transcripts co-expressed with *Bc*TDC-1.

## REFERENCES

1. Wink, M. Ecological Roles of Alkaloids. in Modern Alkaloids: Structure, Isolation, Synthesis, and Biology (eds. Fattorusso, E. & Taglialatela-Scafati, O.) 1–24 (Wiley, 2007). doi:10.1002/9783527621071.

2. Denoeud, F. et al. The coffee genome provides insight into the convergent evolution of caffeine biosynthesis. Science (1979). 345, 1181–1184 (2014).

3. Berman, P. et al. Parallel evolution of cannabinoid biosynthesis. Nat. Plants 9, 817–831 (2023).

4. Chavez, B. G. et al. Elucidation of tropane alkaloid biosynthesis in *Erythroxylum coca* using a microbial pathway discovery platform. Proceedings of the National Academy of Sciences 119, (2022).

5. Pluskal, T. & Weng, J. K. Natural product modulators of human sensations and mood: Molecular mechanisms and therapeutic potential. Chem. Soc. Rev. 47, 1592–1637 (2018).

6. Allen, J. R. F. & Holmstedt, B. R. The simple β-carboline alkaloids. Phytochemistry 19, 157–1582 (1980).

7. Udenfriend, S., Witkop, B., Redfield, B. G. & Weissbach, H. Studies with reversible inhibitors of monoamine oxidase: Harmaline and related compounds. Biochem. Pharmacol. 1, 160–165 (1958).

8. Youdim, M. B. H., Edmondson, D. & Tipton, K. F. The therapeutic potential of monoamine oxidase inhibitors. Nat. Rev. Neurosci. 7, 295–309 (2006).

9. Miller, M. J., Albarracin-Jordan, J., Moore, C. & Capriles, J. M. Chemical evidence for the use of multiple psychotropic plants in a 1,000-year-old ritual bundle from South America. Proceedings of the National Academy of Sciences 116, 11207–11212 (2019).

10. Mckenna, D. J., Towersa, G. H. N. & Abbottb, F. Monoamine oxidase inhibitors in South American hallucinogenic plants: tryptamine and β-carboline constituents of ayahuasca. Journal of Ethnopharmocology 10, 195–223 (1984).

11. Moloudizargari, M., Mikaili, P., Aghajanshakeri, S., Asghari, M. & Shayegh, J. Pharmacological and therapeutic effects of *Peganum harmala* and its main alkaloids. Pharmacogn. Rev. 7, 199–212 (2013).

12. Huber, B. et al. Metabolic profiling reveals first evidence of fumigating drug plant *Peganum harmala* in Iron Age Arabia. *Commun*. Biol. 8, 720 (2025).

13. Shao, H., Huang, X., Zhang, Y. & Zhang, C. Main alkaloids of *Peganum harmala* L. and their different effects on dicot and monocot crops. Molecules 18, 2623–2634 (2013).

14. Liu, Q. et al. Bioactivities and structure–activity relationships of harmine and its derivatives: a review. Chem. Biodivers. 22, e202402953 (2025).

15. Wang, P. et al. A high-throughput chemical screen reveals that harmine-mediated inhibition of DYRK1A increases human pancreatic beta cell replication. Nat. Med. 21, 383–388 (2015).

16. Rosselot, C. et al. Harmine and exendin-4 combination therapy safely expands human β cell mass in vivo in a mouse xenograft system. Sci. Transl. Med. 16, eadg3456 (2024).

17. Stolle, K. & Gröger, D. Untersuchungen zur Biosynthese des Harmins. Arch. Pharm. (Weinheim). 301, 561–571 (1968).

18. Nettleship, L. & Slaytor, M. Limitations of feeding experiments in studying alkaloid biosynthesis in *Peganum harmala* callus cultures. Phytochemistry 13, 735–742 (1974).

19. Davis, C. C. & Choisy, P. Medicinal plants meet modern biodiversity science. Current Biology 34, R158–R173 (2024).

20. Lau, W. & Sattely, E. S. Six enzymes from mayapple that complete the biosynthetic pathway to the etoposide aglycone. Science (1979). 349, 1224–1228 (2015).

21. Nett, R. S., Lau, W. & Sattely, E. S. Discovery and engineering of colchicine alkaloid biosynthesis. Nature 584, 148–153 (2020).

22. Nett, R. S., Dho, Y., Low, Y.-Y. & Sattely, E. S. A metabolic regulon reveals early and late acting enzymes in neuroactive Lycopodium alkaloid biosynthesis. Proceedings of the National Academy of Sciences 118, e2102949118 (2021).

23. Torrens-Spence, M. P. et al. Structural basis for divergent and convergent evolution of catalytic machineries in plant aromatic amino acid decarboxylase proteins. Proceedings of the National Academy of Sciences 117, 10806–10817 (2020).

24. Peyret, H. & Lomonossoff, G. P. The pEAQ vector series: The easy and quick way to produce recombinant proteins in plants. Plant Mol. Biol. 83, 51–58 (2013).

25. Xu, H. et al. Integration of de novo chromosome-level genome and population resequencing of *Peganum* (Nitrariaceae): a case study of speciation and evolutionary trajectories in arid central Asia. Mol. Ecol. Resour. 25, e14078 (2025).

26. Zhao, J. et al. Functional characterization and evolutionary insights into β-carboline *O*-methyltransferases in *Peganum multisectum*. Plant Journal 126, e70998 (2026).

27. Hahn, G. & Ludewig, H. Synthese von Tetrahydro-harman-Derivaten unter physiologischen Bedingungen, I. (vorläuf.) Mitteil. Eur. J. Inorg. Chem. 67, 2031–2035 (1934).

28. Hahn, G., Bärwald, L., Schales, O. & Werner, H. Synthese von tetrahydro-harman (4-carbolin)-derivaten unter physiologischen bedingungen. Justus Liebigs Ann. Chem. 520, 107–123 (1935).

29. Harvey, D. G. & Robson, W. The synthesis of r-6-methoxytryptophan and of harmine, with a note on the action of acetaldehyde on tryptophan. J. Chem. Soc. 97–101 (1938).

30. McIsaac, W. M. Formation of 1-methyl-6-methoxy-1,2,3,4-tetrahydro-2-carboline under physiological conditions. Biochim. Biophys. Acta 52, 607–609 (1961).

31. Callaway, J. C., Gyntber, J., Poso, A., Airaksinen, M. M. & Vepsäläinen, J. The Pictet-Spengler reaction and biogenic tryptamines: Formation of tetrahydro-β-carbolines at physiological pH. J. Heterocycl. Chem. 31, 431–435 (1994).

32. Hester, J. B. Cyclodehydration reactions of tryptamine derivatives with acetone. J. Org. Chem. 29, 2864–2867 (1964).

33. Brandt, S. D., Freeman, S., McGagh, P., Abdul-Halim, N. & Alder, J. F. An analytical perspective on favoured synthetic routes to the psychoactive tryptamines. J. Pharm. Biomed. Anal. 36, 675–691 (2004).

34. Stöckigt, J., Antonchick, A. P., Wu, F. & Waldmann, H. The Pictet-Spengler reaction in nature and in organic chemistry. Angew. Chem. Int. Ed. 50, 8538–8564 (2011).

35. Kürsteiner, O., Dupuis, I. & Kuhlemeier, C. The *pyruvate decarboxylase1* gene of *Arabidopsis* is required during anoxia but not other environmental stresses. Plant Physiol. 132, 968–978 (2003).

36. Evans, S. E., Hu, A. & Phillips, M. A. The plant pyruvate hub. Trends Plant Sci. 31, 238– 250 (2026).

37. Tadege, M. et al. Activation of plant defense responses and sugar efflux by expression of pyruvate decarboxylase in potato leaves. The Plant Journal 16, 661–671 (1998).

38. Yadav, S. K., Singla-Pareek, S. L. & Sopory, S. K. An overview on the role of methylglyoxal and glyoxalases in plants. Drug Metabol. Drug Interact. 23, 51–68 (2008).

39. Schmitz, J., Rossoni, A. W. & Maurino, V. G. Dissecting the physiological function of plant glyoxalase I and glyoxalase I-like proteins. Front. Plant Sci. 871, 1618 (2018).

40. Joshi, V., Laubengayer, K. M., Schauer, N., Fernie, A. R. & Jander, G. Two *Arabidopsis* threonine aldolases are nonredundant and compete with threonine deaminase for a common substrate pool. Plant Cell 18, 3564–3575 (2007).

41. Fujiwara, T. et al. Sekiguchi lesion gene encodes a cytochrome P450 monooxygenase that catalyzes conversion of tryptamine to serotonin in rice. Journal of Biological Chemistry 285, 11308–11313 (2010).

42. Commisso, M. et al. Indolamine accumulation and *TDC/T5H* expression profiles reveal the complex and dynamic regulation of serotonin biosynthesis in tomato (*Solanum lycopersicum* L.). Front. Plant Sci. 13, 975434 (2022).

43. Lombe, B. K., Zhou, T., Caputi, L., Ploss, K. & O’Connor, S. E. Biosynthetic origin of the methoxy group in quinine and related alkaloids. Angew. Chem. Int. Ed. 64, e202418306 (2025).

44. Christensen, J. M. et al. Biosynthesis of pathogen-induced hydroxylated tryptamine derivatives in barley. Plant Physiol. 202, kiag622 (2026).

45. Arnao, M. B. Phytomelatonin: discovery, content, and role in plants. Advances in Botany 2014, 815769 (2014).

46. Erland, L. A. E., Murch, S. J., Reiter, R. J. & Saxena, P. K. A new balancing act: The many roles of melatonin and serotonin in plant growth and development. Plant Signal. Behav. 10, e1096469 (2015).

47. Strommer, J. The plant *ADH* gene family. The Plant Journal 66, 128–142 (2011).

48. Havir, E. A. & McHale, N. A. Biochemical and developmental characterization of multiple forms of catalase in tobacco leaves. Plant Physiol. 84, 450–455 (1987).

49. Mehta, N., Meng, Y., Zare, R., Kamenetsky-Goldstein, R. & Sattely, E. A developmental gradient reveals biosynthetic pathways to eukaryotic toxins in monocot geophytes. Cell 187, 5620–5637.e10 (2024).

50. Byeon, H.-J., Jung, K.-H., Moon, G.-S., Moon, S.-K. & Lee, H.-Y. A facile and efficient method for the synthesis of crystalline tetrahydro-β-carbolines via the Pictet-Spengler reaction in water. Sci. Rep. 10, 1057 (2020).

51. Cai, L. et al. Deeply altered genome architecture in the endoparasitic flowering plant *Sapria himalayana* Griff. (Rafflesiaceae). Current Biology 31, 1002–1011.e9 (2021).

52. Cheng, H., Concepcion, G. T., Feng, X., Zhang, H. & Li, H. Haplotype-resolved de novo assembly using phased assembly graphs with hifiasm. Nat. Methods 18, 170–175 (2021).

53. Gurevich, A., Saveliev, V., Vyahhi, N. & Tesler, G. QUAST: quality assessment tool for genome assemblies. Bioinformatics 29, 1072–1075 (2013).

54. Rhie, A., Walenz, B. P., Koren, S. & Phillippy, A. M. Merqury: reference-free quality, completeness, and phasing assessment for genome assemblies. Genome Biol. 21, 245 (2020).

55. Storer, J., Hubley, R., Rosen, J., Wheeler, T. J. & Smit, A. F. The Dfam community resource of transposable element families, sequence models, and genome annotations. Mob. DNA 12, 2 (2021).

56. Flynn, J. M. et al. RepeatModeler2 for automated genomic discovery of transposable element families. Proceedings of the National Academy of Sciences 117, 9451–9457 (2020).

57. Chen, N. Using RepeatMasker to identify repetitive elements in genomic sequences. Curr. Protoc. Bioinformatics 5, 4–10 (2004).

58. Freedman, A. H. & Sackton, T. B. Building better genome annotations across the tree of life. Genome Res. 35, 1261–1276 (2025).

59. Pertea, M. et al. StringTie enables improved reconstruction of a transcriptome from RNA-seq reads. Nat. Biotechnol. 33, 290–295 (2015).

60. Fu, L., Niu, B., Zhu, Z., Wu, S. & Li, W. CD-HIT: Accelerated for clustering the next-generation sequencing data. Bioinformatics 28, 3150–3152 (2012).

61. Bolger, A. M., Lohse, M. & Usadel, B. Trimmomatic: A flexible trimmer for Illumina sequence data. Bioinformatics 30, 2114–2120 (2014).

62. Bray, N. L., Pimentel, H., Melsted, P. & Pachter, L. Near-optimal probabilistic RNA-seq quantification. Nat. Biotechnol. 34, 525–527 (2016).

63. Haas, B. J. et al. De novo transcript sequence reconstruction from RNA-seq using the Trinity platform for reference generation and analysis. Nat. Protoc. 8, 1494–1512 (2013).

64. Eddy, S. R. Accelerated profile HMM searches. PLoS Comput. Biol. 7, e1002195 (2011).

65. Camacho, C. et al. BLAST+: architecture and applications. BMC Bioinformatics 10, 421 (2009).

66. Armenteros, J. J. A. et al. Detecting sequence signals in targeting peptides using deep learning. Life Sci. Alliance 2, e201900429 (2019).

67. Sperschneider, J. et al. LOCALIZER: subcellular localization prediction of both plant and effector proteins in the plant cell. Sci. Rep. 7, 44598 (2017).

68. Stoddard, A. & Rolland, V. I see the light! Fluorescent proteins suitable for cell wall/apoplast targeting in *Nicotiana benthamiana* leaves. Plant Direct 3, e00112 (2019).

69. Schindelin, J. et al. Fiji: an open-source platform for biological-image analysis. Nat. Methods 9, 676–682 (2012).

70. Katoh, K. & Standley, D. M. MAFFT multiple sequence alignment software version 7: improvements in performance and usability. Mol. Biol. Evol. 30, 772–780 (2013).

71. Gouet, P., Courcelle, E., Stuart, D. I. & Métoz, F. ESPript: analysis of multiple sequence alignments in PostScript. Bioinformatics 15, 305–308 (1999).

72. Minh, B. Q. et al. IQ-TREE 2: new models and efficient methods for phylogenetic inference in the genomic era. Mol. Biol. Evol. 37, 1530–1534 (2020).

73. Kalyaanamoorthy, S., Minh, B. Q., Wong, T. K. F., von Haeseler, A. & Jermiin, L. S. ModelFinder: fast model selection for accurate phylogenetic estimates. Nat. Methods 14, 587–589 (2017).

74. Hoang, D. T., Chernomor, O., von Haeseler, A., Minh, B. Q. & Vinh, L. S. UFBoot2: improving the ultrafast bootstrap approximation. Mol. Biol. Evol. 35, 518–522 (2018).

75. Shimodaira, H. An approximately unbiased test of phylogenetic tree selection. Syst. Biol. 51, 492–508 (2002).

76. Letunic, I. & Bork, P. Interactive Tree of Life (iTOL) v6: recent updates to the phylogenetic tree display and annotation tool. Nucleic Acids Res. 52, W78–W82 (2024).

