## Supplementary Information for "Independent evolution of harmala alkaloids in medicinal Syrian rue and ayahuasca plants"

**page(s)**

**Supplementary Figures**........................................3-33

Figure 1. Harmala alkaloid quantification in *Peganum harmala* seedling tissues via LC-MS .......... 3
Figure 2. Distribution of $D_5$-tryptophan ($D_5$-trp) and labeled harmala alkaloid distribution in *P. harmala* seedlings fed with $D_5$-trp .......... 4
Figure 3. Tissue-specific expression of pyridoxal phosphate (PLP)-dependent decarboxylase genes in *P. harmala* seedlings and characterization of tryptophan decarboxylases (TDCs) .......... 5
Figure 4. Stepwise reconstitution of harmine (6) biosynthesis from *P. harmala* via transient expression in *N. benthamiana* .......... 6
Figure 5. *Ph*OMT-1 substrate specificity .......... 7
Figure 6. *Ph*FDO-1 and *Ph*FDO-2 substrate specificity .......... 8
Figure 7. Tandem array of FDO genes in *Peganum multisectum* .......... 9
Figure 8. Spontaneous scaffold formation by 6′-hydroxytryptamine (2) condensation with pyruvate. .......... 10-11
Figure 9. Spontaneous scaffold formation by 6′-hydroxytryptamine (2) condensation with acetaldehyde. .......... 12
Figure 10. Reconstituting the harmala alkaloid biosynthetic pathway from *P. harmala* with native or truncated *Ph*PDC .......... 13
Figure 11. Identification of an N-terminal plastid localization signal in *Ph*PDC-1 .......... 14
Figure 12. Harmine (6) biosynthesis in *N. benthamiana* transiently expressing *P. harmala* genes and a pyruvate decarboxylase from *Zymomonas mobilis* (*Zm*PDC) .......... 15
Figure 13. Phylogenetic and active site sequence analysis of *Ph*GLXI-like .......... 16
Figure 14. Effect of *Ph*GLXI-like on harmala alkaloid biosynthesis .......... 17
Figure 15. Identification of a tryptophan decarboxylase from *B. caapi* .......... 18
Figure 16. *Bc*OMT-1 and *Bc*OMT-2 substrate specificity .......... 19
Figure 17. *Bc*FDO-1 substrate specificity .......... 20
Figure 18. Tetrahydroharmine (8) biosynthesis in *P. harmala* and *B. caapi* using transient expression in *N. benthamiana* .......... 21-22
Figure 19. Maximum likelihood phylogenetic tree of threonine aldolases (TAs) analyzed in this study. .......... 23

Figure 20. Maximum likelihood phylogenetic tree of tryptamine hydroxylases..................................... 24
Figure 21. Characterization of putative tryptophan 5'-hydroxylases (T5Hs) from diverse plant orders.. 25
Figure 22. Protein multiple sequence alignment of characterized tryptamine hydroxylases .............26-28
Figure 23. Comparison of predicted T6H and T5H structures from *P. harmala* and *B. caapi*................ 29
Figure 24. Characterization of mutagenized T6H enzymes from *P. harmala* and *B. caapi* .................... 30
Figure 25. Maximum likelihood phylogenetic tree of *O*-methyltransferases (OMTs) evaluated in this study ........................................................................................................................ 31
Figure 26. Maximum likelihood phylogenetic tree of flavin-dependent oxidases (FDOs) analyzed in this study ........................................................................................................................ 32
Figure 27. Summary of harmala alkaloid biosynthesis in *P. harmala* and *B. caapi*................................ 33

**Supplementary Methods** ........................................................................................................ 34

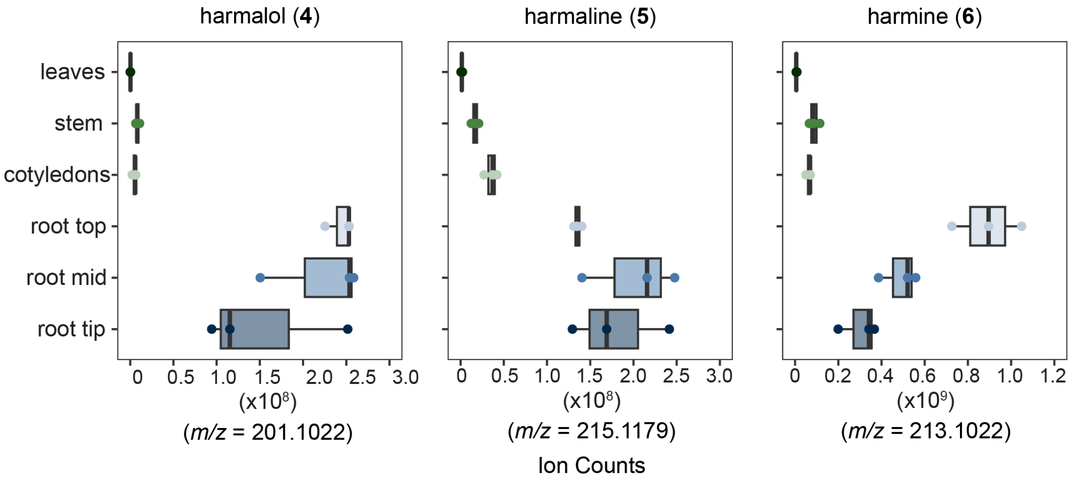

**SI Fig. 1. Harmala alkaloid quantification in *Peganum harmala* seedling tissues via LC-MS.** Dots represent biological replicates, and box plots depict the median, first, and third quartiles ($n$ = 3).

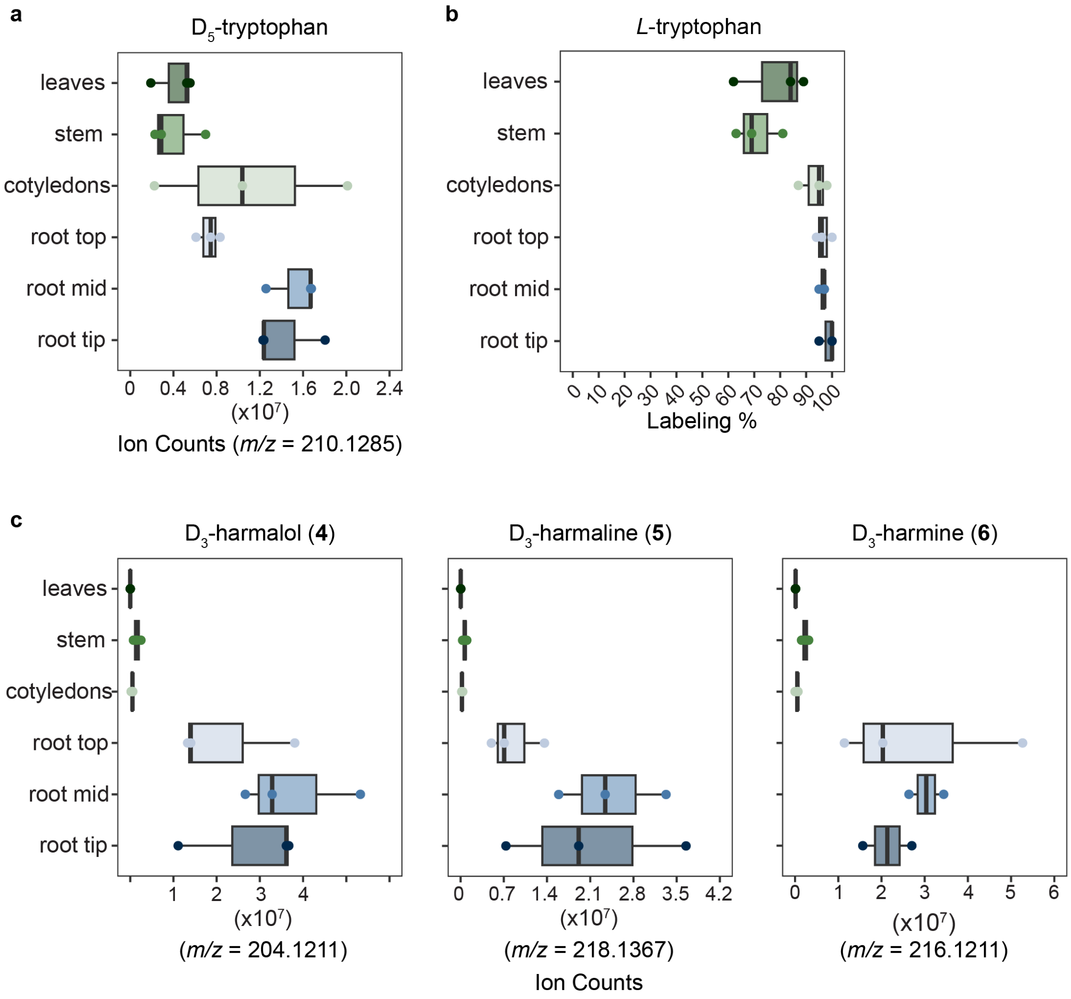

**SI Fig. 2. Distribution of $D_5$-tryptophan ($D_5$-trp) and labeled harmala alkaloid distribution in *P. harmala* seedlings fed with $D_5$-trp. a)** $D_5$-trp distribution across tissues. **b)** Labeling percentage of tryptophan calculated by dividing labeled tryptophan by the sum of labeled and unlabeled tryptophan. **c)** Distribution of labeled harmalol (**4**), harmaline (**5**), and harmine (**6**). Data for harmine also shown in Figures 2b. Dots represent biological replicates, and box plots depict the median, first, and third quartiles (*n* = 3).

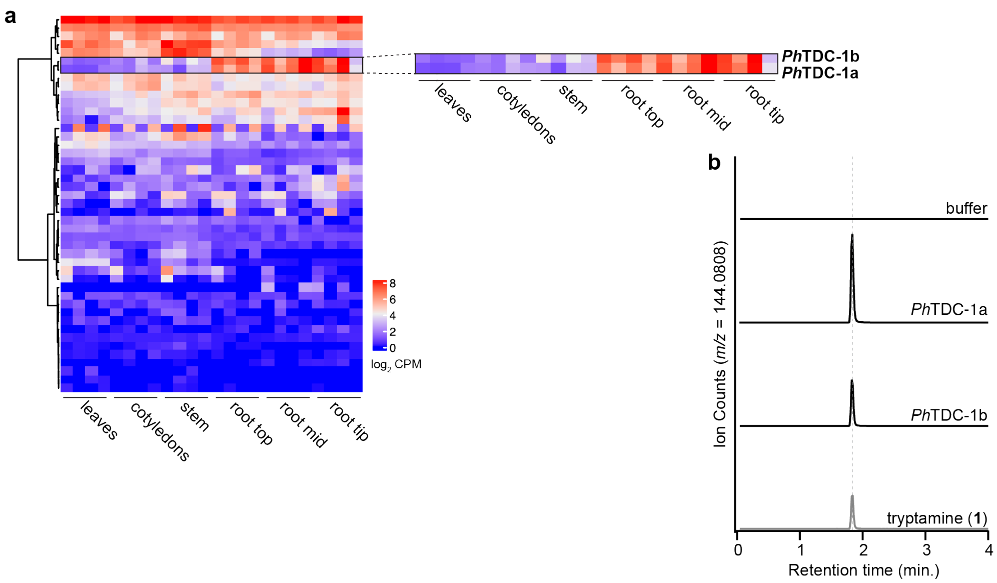

**SI Fig. 3. Tissue-specific expression of pyridoxal phosphate (PLP)-dependent decarboxylase genes in *P. harmala* seedlings and characterization of tryptophan decarboxylases (TDCs). a)** Gene expression heatmap of the 45 transcripts annotated as PLP-dependent decarboxylases in the *P. harmala* transcriptome. Columns represent four biological replicates of each tissue type and rows are the $log_2$-transformed counts per million (CPM) clustered by Euclidean distance based on the ward.D method. Highlighted are two putative TDC genes. **b)** Characterization of putative TDCs by transient expression in *N. benthamiana* and metabolite analysis by LC-MS with a tryptamine (**1**) standard. The buffer control is for a leaf infiltrated with Agrobacterium induction media. Ion counts represent the major in-source fragment detected by LC-MS ($[M\text{-}NH_3+H]^+$ = *m/z* 144.0808).

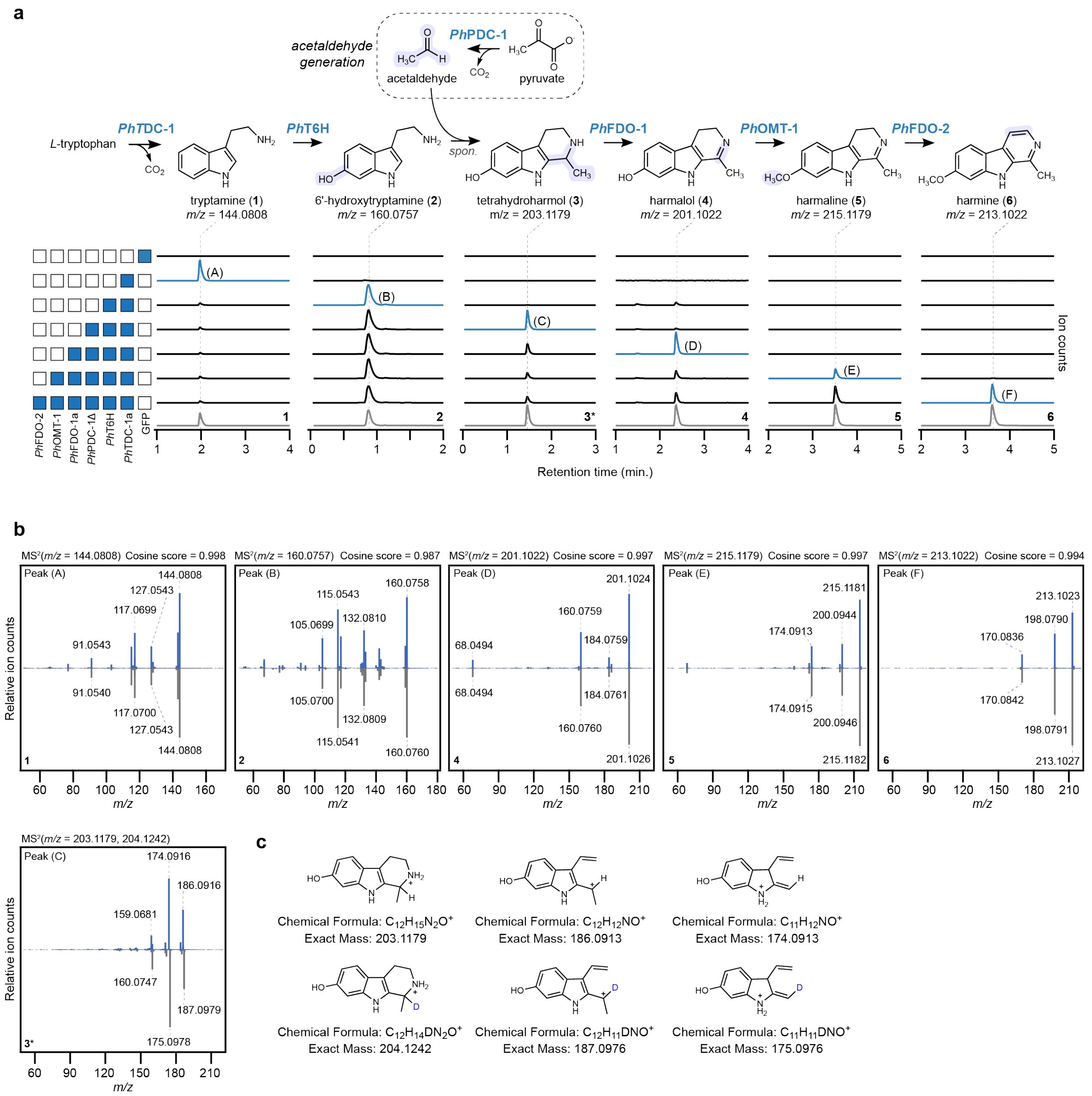

**SI Fig 4. Stepwise reconstitution of harmine (6) biosynthesis from *P. harmala* via transient expression in *N. benthamiana*.** **a**) Shown are EICs for the mass ion (*m/z*) of each intermediate compared to a standard (gray). Blue boxes to the left of bar graphs indicate inclusion of a gene in transient expression experiments, while white boxes indicate omission of a gene. Traces in blue are the product of the minimal set of enzymes required to produce a given intermediate (A-F). A deuterated standard was used for tetrahydroharmol (**3**)* ($D_2$-tetrahydroharmol) $[M+H]^+$ = *m/z* 204.1242). spon. = spontaneous. **b**) MS/MS fragmentation spectra (20V) of each intermediate (blue) compared to an authentic standard (gray). The most abundant ion for each compound was selected for fragmentation, and it is indicated above each spectrum. **c**) Proposed structures for MS/MS ions observed for peak (C) and **3***.

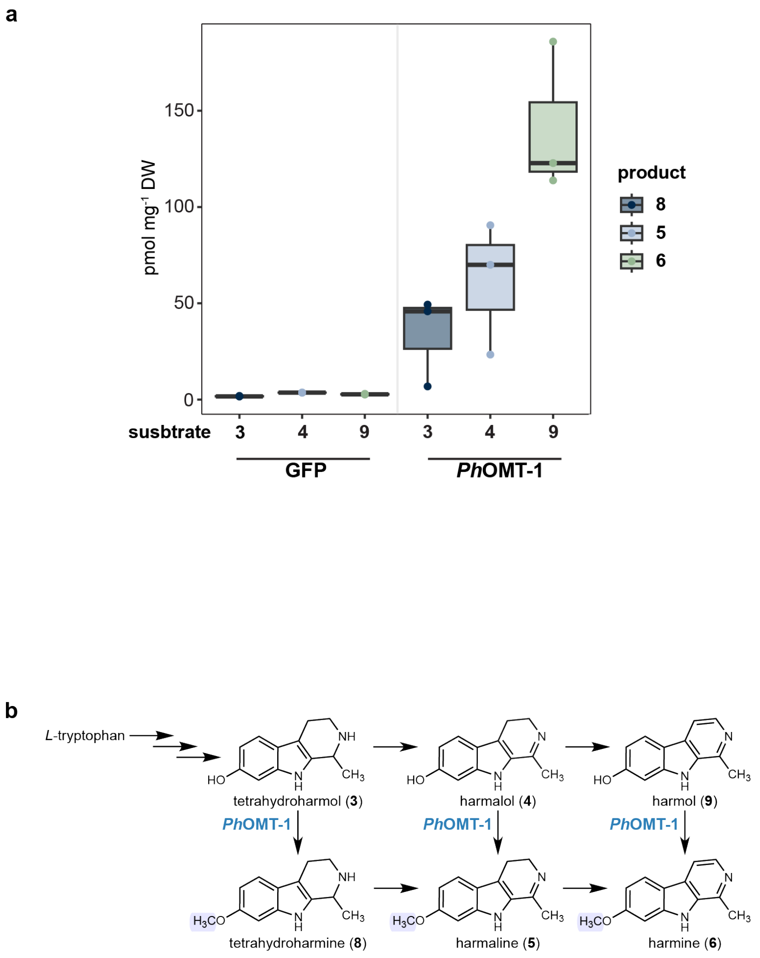

**SI Fig. 5. *Ph*OMT-1 substrate specificity. a)** *Ph*OMT-1 and GFP (control) were transiently expressed in *N. benthamiana* leaves and infiltrated with 50 µM of substrate ($D_2$-tetrahydroharmol (**3**), harmalol (**4**), or harmol (**9**)). The corresponding *O*-methylated products ($D_2$-tetrahydroharmine (**8**), harmaline (**5**), and harmine (**6**)) were quantified from plant methanolic extracts by LC-MS using standard curves. Dots represent biological replicates, and box plots depict the median, first, and third quartiles ($n = 3$). **b)** Metabolic network of harmala alkaloids depicting reactions catalyzed by *Ph*OMT-1.

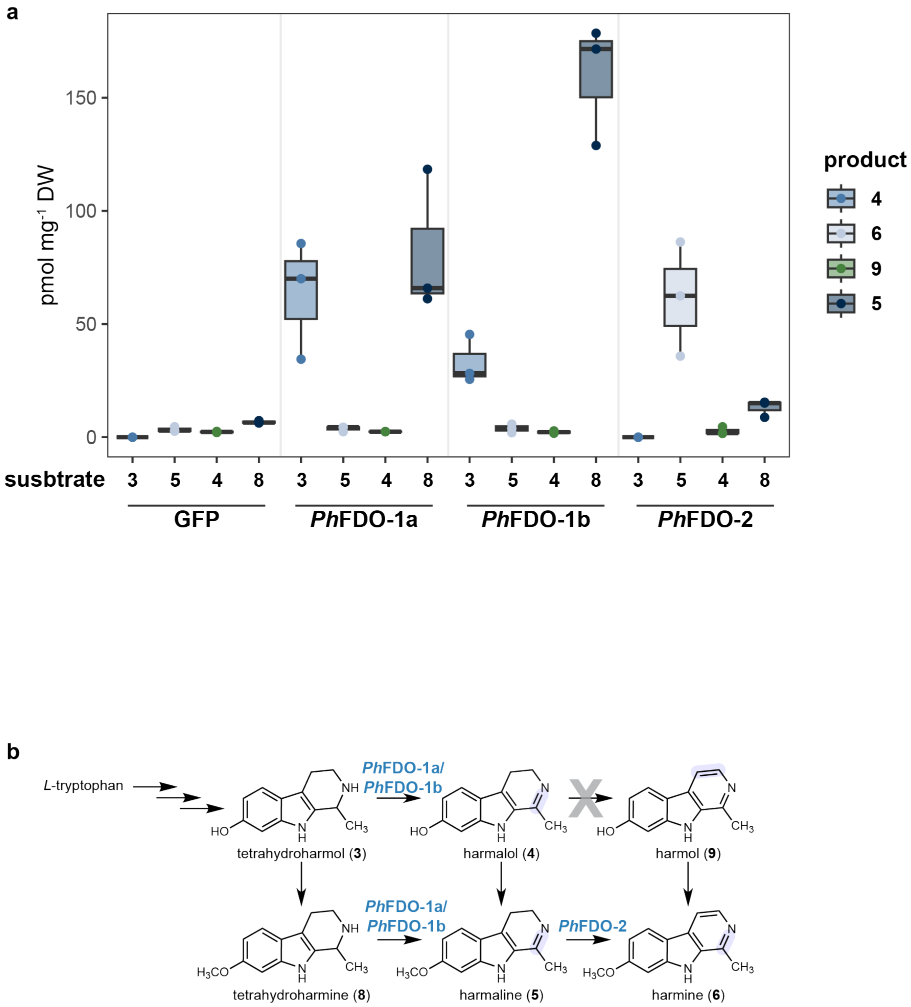

**SI Fig. 6. *Ph*FDO-1 and *Ph*FDO-2 substrate specificity. a)** *Ph*FDO-1a, *Ph*FDO-1b, *Ph*FDO-2, and GFP (control) were transiently expressed in *N. benthamiana* leaves and infiltrated with 50 µM of substrate ($D_2$-tetrahydroharmol (**3**), harmaline (**5**), harmalol (**4**), or tetrahydroharmine (**8**)). The corresponding oxidized products (harmalol (**4**), harmine (**6**), harmol (**9**), and harmaline (**5**)) were quantified from plant methanolic extracts by LC-MS using standard curves. Dots represent biological replicates, and box plots depict the median, first, and third quartiles ($n = 3$). **b)** Metabolic network of harmala alkaloids depicting reactions catalyzed by *Ph*FDO-1a, *Ph*FDO-1b, *Ph*FDO-2.

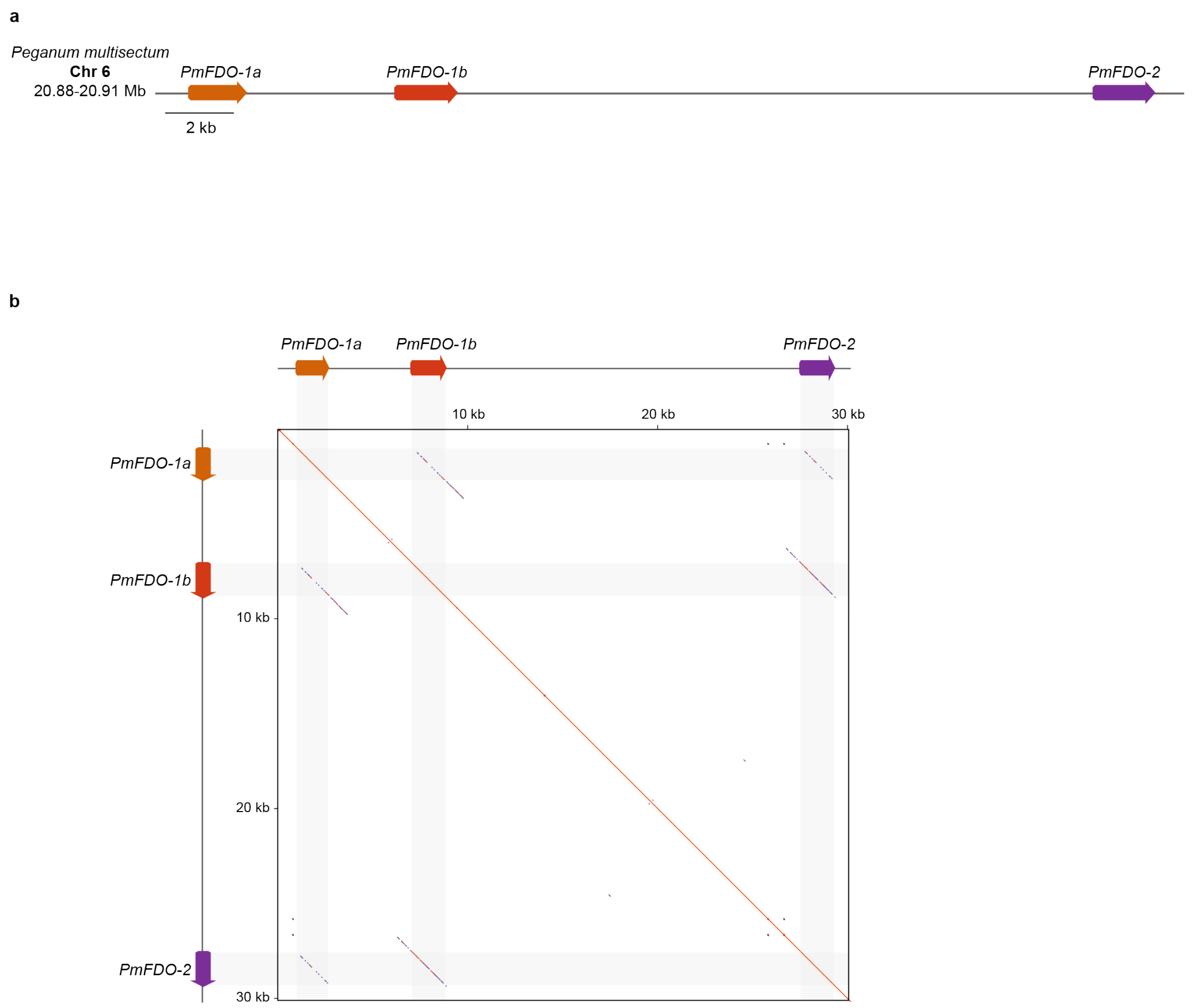

**SI Fig 7. Tandem array of FDO genes in *Peganum multisectum*. a)** Genomic locus in *P. multisectum* containing orthologs of the *P. harmala FDO-1a*, *FDO-1b*, and *FDO-2* genes within a ~30 kilobase (kb) region. **b)** Dot plot (word size = 20) representing the sequence similarity among FDO genes in this locus.

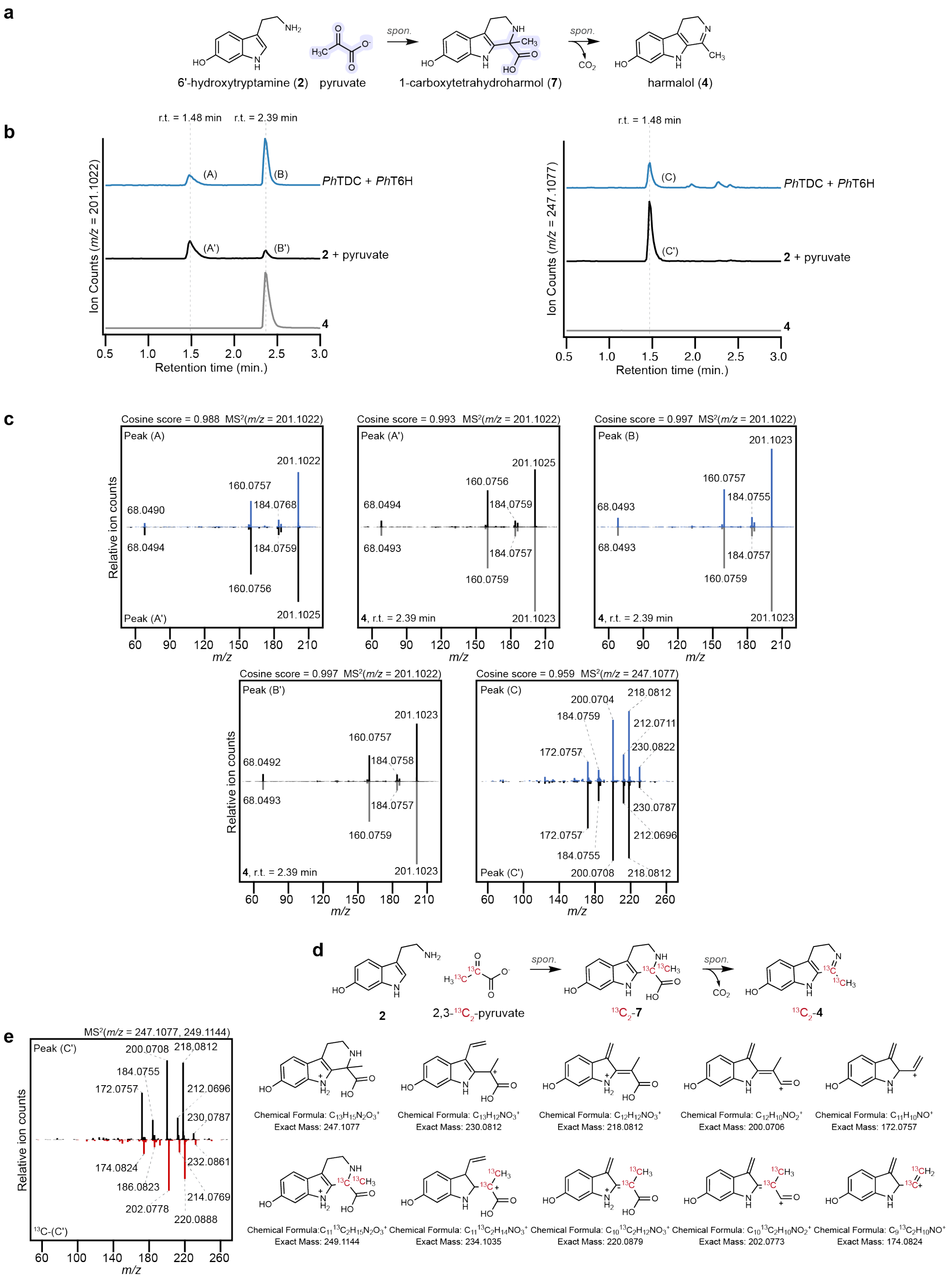

**SI Fig 8. Spontaneous scaffold formation by 6′-hydroxytryptamine (2) condensation with pyruvate. a**) Diagram of Pictet-Spengler condensation of 6′-hydroxytryptamine (**2**) and pyruvate to form 1-carboxytetrahydroharmol (**7**) and spontaneous oxidative decarboxylation of (**7**) to form harmalol (**4**). **b**) EICs for the reaction products depicted in panel (a) generated through transient gene expression in *N. benthamiana* (blue) and via *in vitro* (black) reactions with (**2**) and pyruvate under acidic buffer conditions. On the left chromatogram, peaks A and A′ (1.48 min) represent a proposed in-course ion fragment resulting from the oxidative decarboxylation of (**7**). Peaks B and B′ (2.39 min) are confirmed to be (**4**) via comparison to an authentic standard. On the right chromatogram, peaks C and C′ (1.48 min) represent the $[M+H]^+$ ion for (**7**). **c**) MS/MS fragmentation spectra (20V) of peaks described in panel (b). Note that the MS/MS spectrum for the in-source fragment of peak A is identical to the spectrum for (**4**), supporting the proposed in-source oxidative decarboxylation of (**7**) during MS analysis. **d**) Diagram for the condensation of (**2**) and 2,3-$^{13}C_2$-pyruvate to form $^{13}C_2$-carboxytetrahydroharmol $^{13}C_2$-(**7**) under acidic buffer conditions. **e**) MS/MS spectrum of (**7**) and $^{13}C_2$-(**7**) formed *in vitro* (left) and proposed structures for the observed ion fragments (right).

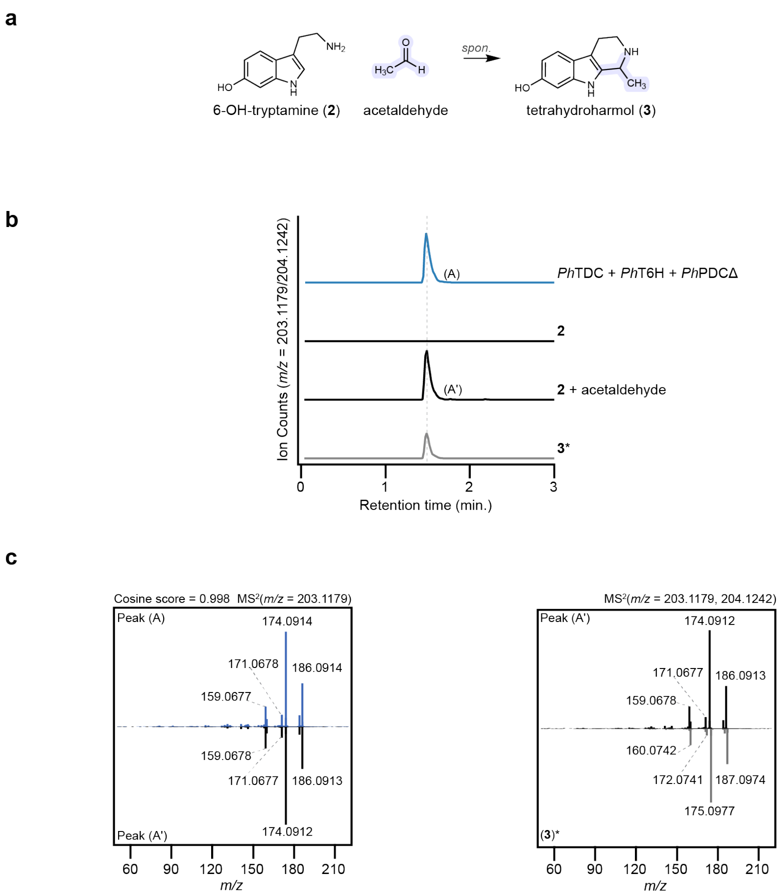

**SI Fig 9. Spontaneous scaffold formation by 6′-hydroxytryptamine (2) condensation with acetaldehyde.** **a**) Diagram of Pictet-Spengler condensation of 6′-hydroxytryptamine (**2**) and acetaldehyde to form tetrahydroharmol (**3**). **b**) EICs for the reaction product depicted in panel (a) generated through transient gene expression in *N. benthamiana* (blue) and via *in vitro* (black) reactions with (**2**) and acetaldehyde under acidic buffer conditions. **c**) MS/MS fragmentation spectra (20V) of peaks described in panel (b).

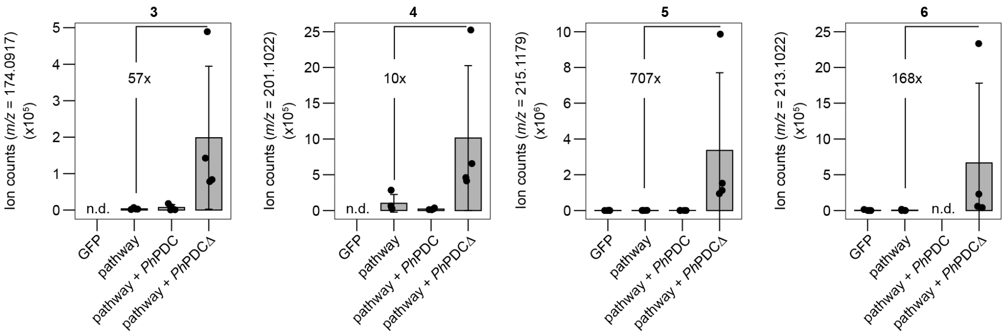

**SI Fig 10. Reconstituting the harmala alkaloid biosynthetic pathway from *P. harmala* with native or truncated *Ph*PDC.** Shown are quantified EICs for the mass ion (*m/z*) of tetrahydroharmol (**3**), harmalol (**4**), harmaline (**5**), and harmine (**6**) produced by *N. benthamiana* leaves transiently expressing GFP (control), pathway (*Ph*TDC-1a, *Ph*T6H, *Ph*OMT-1, *Ph*FDO-1b, and *Ph*FDO-2), pathway with native *Ph*PDC, and pathway with *Ph*PDCΔ (N-terminal signal peptide removed). For tetrahydroharmol (**3**), an in-source fragment ($[M\text{-}CNH_3\text{+}H]^+$ = *m/z* 174.0917) was used for quantifying ion abundance. Dots represent biological replicates ($n$ = 4) and bar graphs represent the mean ± s.d. Mean fold change in between leaves expressing pathway and pathway with *Ph*PDCΔ is shown for each intermediate. Samples were compared by one-way ANOVA ($p > 0.05$ among strain combinations for each intermediate shown here). n.d. = not detected

a

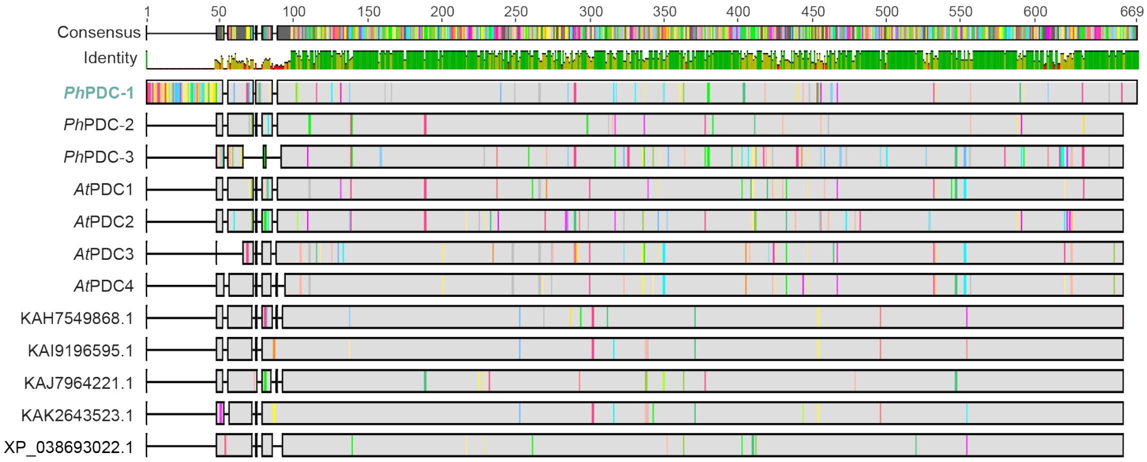

b

| Protein_ID | Localizations | Plastid | Cytoplasm |
|---|---|---|---|
| PhPDC-1 | Plastid | 0.97 | 0.16 |
| PhPDC-2 | Cytoplasm | 0.05 | 0.67 |
| PhPDC-3 | Cytoplasm | 0.12 | 0.65 |
| AtPDC1 | Cytoplasm | 0.06 | 0.66 |
| AtPDC2 | Cytoplasm | 0.06 | 0.67 |
| AtPDC3 | Cytoplasm | 0.11 | 0.66 |
| AtPDC4 | Cytoplasm | 0.10 | 0.69 |
| KAH7549868.1 | Cytoplasm | 0.04 | 0.66 |
| KAI9196595.1 | Cytoplasm | 0.05 | 0.67 |
| KAJ7964221.1 | Cytoplasm | 0.06 | 0.63 |
| KAK2643523.1 | Cytoplasm | 0.05 | 0.66 |
| XP_038693022.1 | Cytoplasm | 0.07 | 0.66 |

**SI Fig 11. Identification of an N-terminal plastid localization signal in *Ph*PDC-1**. **a)** Protein sequence alignment (MUSCLE) between *Ph*PDC-1, other PDCs predicted in *P. harmala*, PDC homologs from *Arabidopsis thaliana*, and the closest BLASTP hits from the NCBI non-redundant protein database. The visual for this alignment was generated in Geneious Pro. **b)** Output summary from signal peptide prediction of PDC protein sequences with DeepLoc 2.1 (https://services.healthtech.dtu.dk/services/DeepLoc-2.1/). Numerical values represent probabilities of localization to a particular cellular compartment.

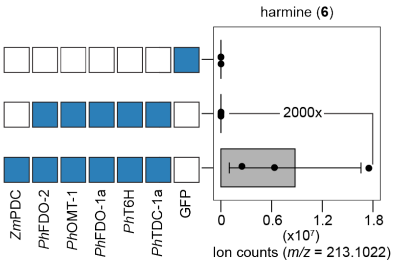

**SI Fig. 12. Harmine (6) biosynthesis in *N. benthamiana* transiently expressing *P. harmala* genes and a pyruvate decarboxylase from *Zymomonas mobilis* (*Zm*PDC).** Leaves were infiltrated with combinations of biosynthetic genes or GFP as a negative control, and harmine (**6**) ion abundance was quantified via LC-MS. Dots represent biological replicates ($n$ = 3) and bar graphs represent the mean ± s.d. Samples were compared by one-way ANOVA ($p$ = 0.08). The 2000x represents the fold change in harmine accumulation in between leaves expressing pathway genes with and without *Zm*PDC.

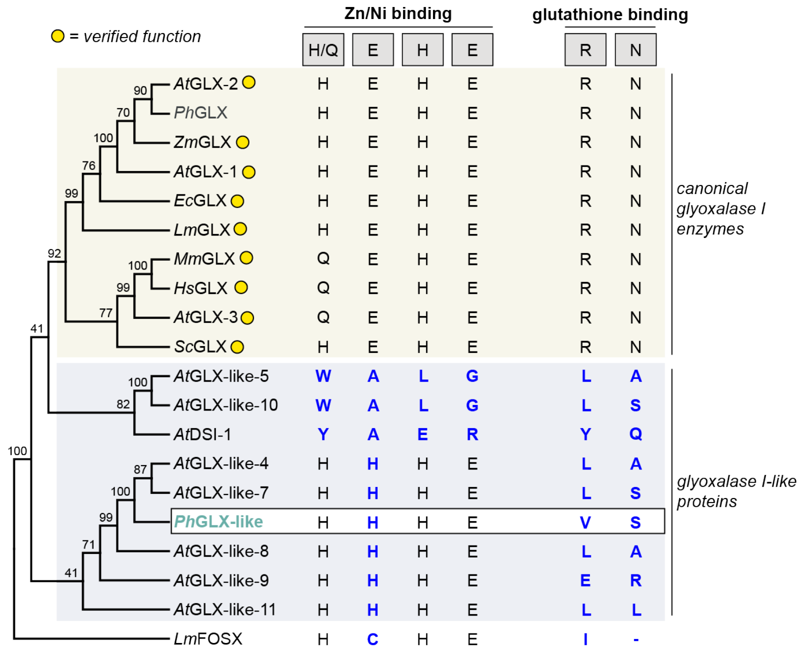

**SI Fig 13. Phylogenetic and active site sequence analysis of *Ph*GLXI-like.** Shown is a phylogenetic tree (MUSCLE alignment, Neighbor-Joining consensus tree using the Jukes-Cantor gene distance model, 1000 bootstrap replicates) containing verified, canonical glyoxalase (GLX) enzymes (indicated with a yellow circle), as well as GLX-like proteins from *Arabidopsis thaliana* and the newly identified *Ph*GLXI-like from Syrian rue. Next to each protein annotation are the amino acids that correspond to the canonical metal and glutathione binding sites in the enzyme active site. Amino acids that differ from the canonical sequence are colored in blue. FosX from *Listeria monocytogenes* (UniProt Q8Y6I2) was used as an outgroup, as it belongs to the same Glyoxalase/Bleomycin resistance protein/Dihydroxybiphenyl dioxygenase superfamily (InterPro IPR029068).

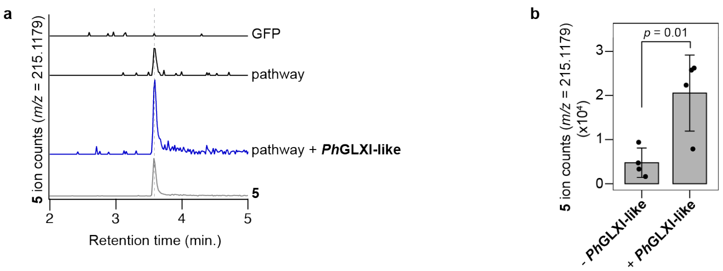

**SI Fig 14. Effect of *Ph*GLXI-like on HA biosynthesis. a)** Harmaline (**5**) biosynthesis in *N. benthamiana* leaves transiently expressing *Ph*TDC-1a, *Ph*T6H, *Ph*OMT-1, *Ph*FDO-1b, and *Ph*FDO-2 (pathway) with and without *Ph*GLXI-like. **b**) Harmaline ion counts in *N. benthamiana* leaves expressing *P. harmala* pathway genes with and without *Ph*GLXI-like. Dots represent biological replicates ($n$ = 4) and bar graphs represent the mean ± s.d. Samples were compared using a two-sided Student's $t$-test, and the resulting $p$-value is displayed above the graph.

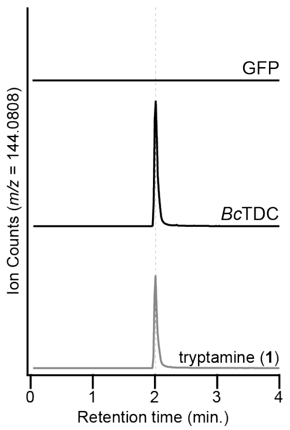

**SI Fig. 15. Identification of a tryptophan decarboxylase from *B. caapi*.** A putative tryptophan decarboxylase (TDC) from *B. caapi* was characterized by transient expression in *N. benthamiana* and metabolite analysis by LC-MS with comparison to a tryptamine (**1**) standard. Shown are EICs for an in-source fragment of tryptamine (**1**) ($[M\text{-}NH_3\text{+}H]^+$ = *m/z* 144.0808). The negative control was an *N. benthamiana* leaf transiently expressing GFP.

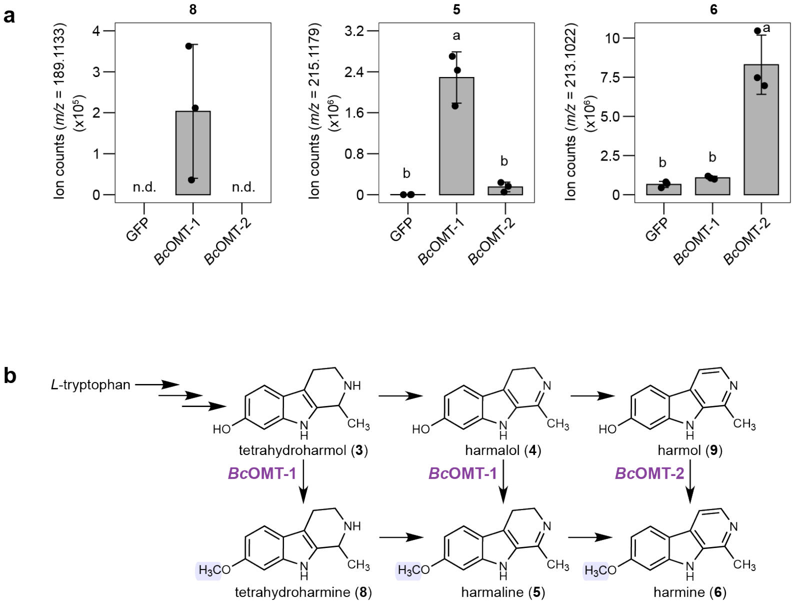

**SI Fig. 16. *Bc*OMT-1 and *Bc*OMT-2 substrate specificity. a)** *Bc*OMT-1, *Bc*OMT-2 and GFP (control) were transiently expressed in *N. benthamiana* leaves and infiltrated with substrate ($D_2$-tetrahydroharmol (**3**), harmalol (**4**), or harmol (**9**)). Shown are the integrated EICs for the corresponding *O*-methylated products ($D_2$-tetrahydroharmine (**8**), harmaline (**5**), and harmine (**6**)). For $D_2$-tetrahydroharmine (**8**), an in-source fragment ($[M\text{-}CNH_3\text{+}H]^+$ = *m/z* 189.1133) was used for quantifying ion abundance. Dots represent biological replicates ($n = 3$) and bar graphs represent the mean ± s.d. The letters above the bar graphs represent differences between strain combinations determined by one-way ANOVA and Tukey's honest significant difference test ($p < 0.05$). n.d. = not detected **b)** Metabolic network of harmala alkaloids depicting reactions catalyzed by *Bc*OMT-1 and *Bc*OMT-2.

a

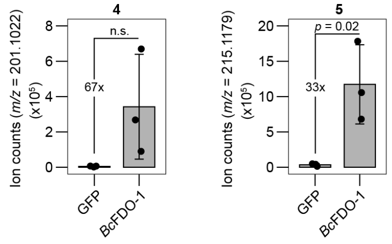

b

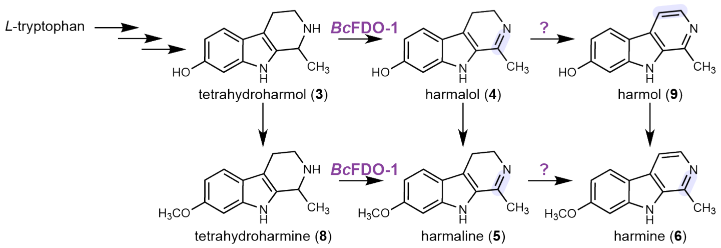

**SI Fig. 17. *Bc*FDO-1 substrate specificity. a)** *Bc*FDO-1 and GFP (control) were transiently expressed in *N. benthamiana* leaves and infiltrated with either $D_2$-tetrahydroharmol (**3**), or tetrahydroharmine (**8**). Shown are the integrated EICs for the corresponding desaturated products, harmalol (**4**) and harmaline (**5**). Dots represent biological replicates ($n$ = 3) and bar graphs represent the mean ± s.d. Samples were compared using a two-sided Student's $t$-test, and the resulting $p$-value is displayed above the graph. n.s. = not significant. **b)** Metabolic network of harmala alkaloids depicting reactions catalyzed by *Bc*FDO-1.

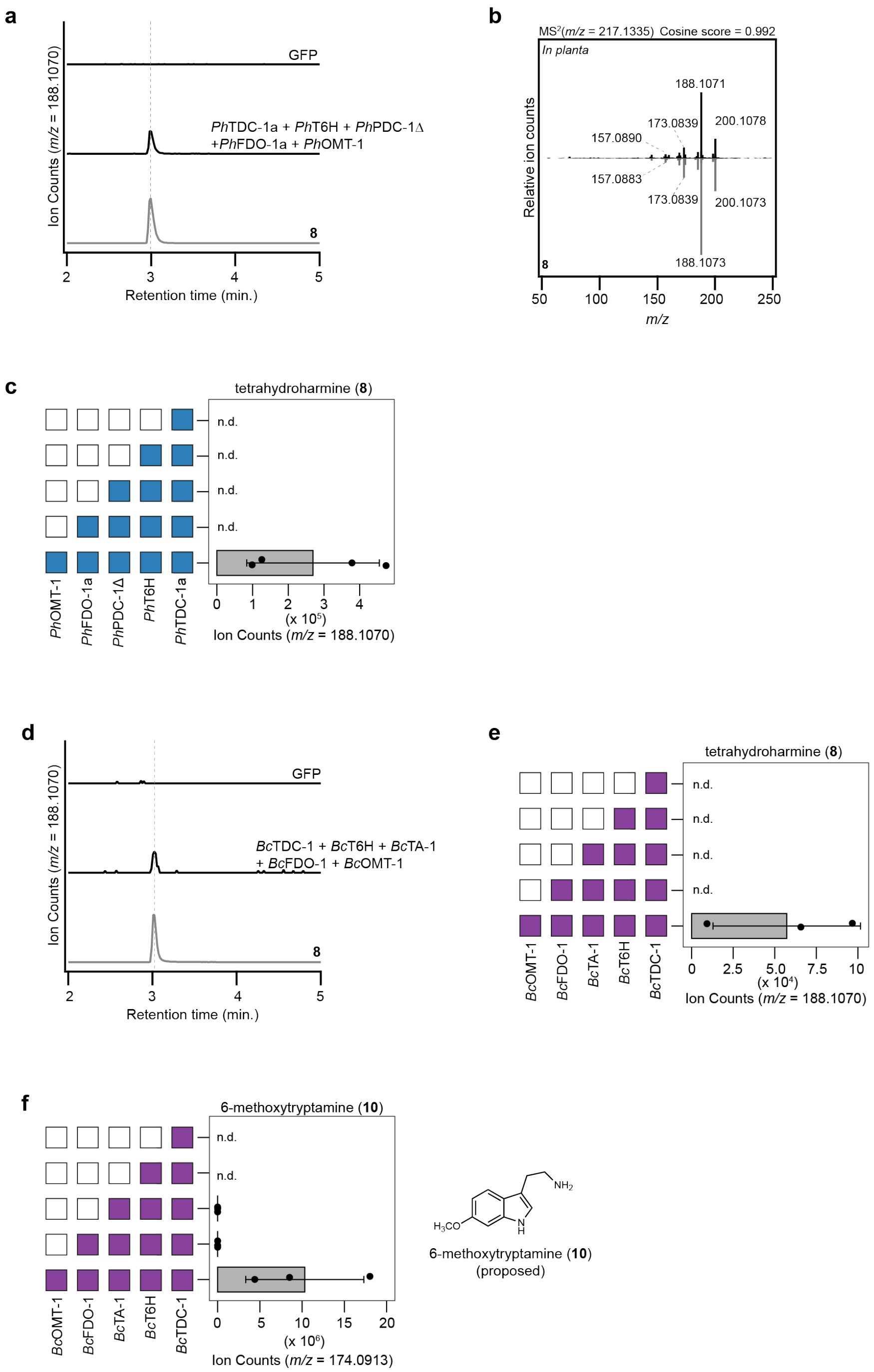

**SI Fig. 18. Tetrahydroharmine (8) biosynthesis in *P. harmala* and *B. caapi* using transient expression in *N. benthamiana*.** **a**) EIC traces for an in-source fragment of (**8**) ($[M\text{-}CNH_3\text{+}H]^+$ = *m/z* 188.1070) in *N. benthamiana* transiently expressing either GFP (negative control) or *P. harmala* genes compared to an authentic standard (gray). **b**) MS/MS spectra of (**8**) produced in *N. benthamiana* by *P. harmala* enzymes (black) compared and an authentic standard (black). **c**) Stepwise reconstitution of (**8**) in *N. benthamiana* using *P. harmala* genes. Dots represent biological replicates ($n$ = 4) and bar graphs represent the mean ± s.d. Blue boxes to the left of bar graphs indicate inclusion of a gene in transient expression experiments, while white boxes indicate omission of a gene. **d**) EIC traces for (**8**) produced by *N. benthamiana* transiently expressing either GFP (negative control) or *B. caapi* genes compared to an authentic standard (gray). **e**) Stepwise reconstitution of (**8**) by transient expression of *B. caapi* genes in *N. benthamiana*. **f**) Production of putative 6-methoxytryptamine ($[M\text{-}NH_3+H]^+$ = *m/z* 174.0913) during transient expression of *B. caapi* genes in *N. benthamiana*. For panels (e) and (f), purple boxes to the left of bar graphs indicate inclusion of a gene in transient expression experiments, while white boxes indicate omission of a gene. n.d. = not detected. Dots represent biological replicates (*n* = 3) and bar graphs represent the mean ± s.d.

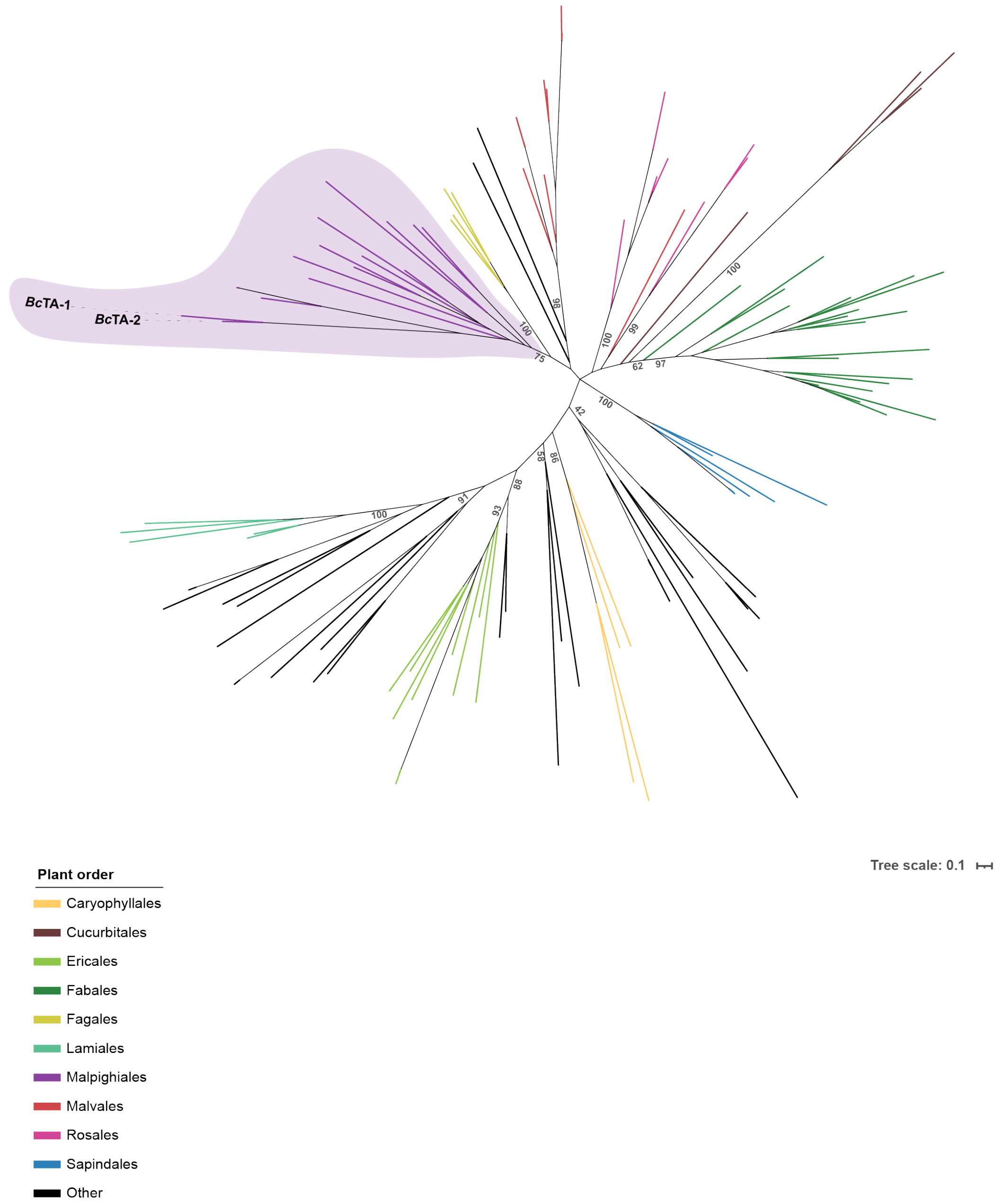

**SI Fig. 19. Maximum likelihood phylogenetic tree of threonine aldolases (TAs) analyzed in this study.** Labeled branches represent biochemically characterized enzymes. Branches are colored by plant phylogenetic order. Branch support for main nodes is indicated at the tree nodes by ultrafast bootstrap approximation (UFBoot) values using 1000 bootstrap resamplings. A complete description of sequences and parameters used can be found in the Methods section.

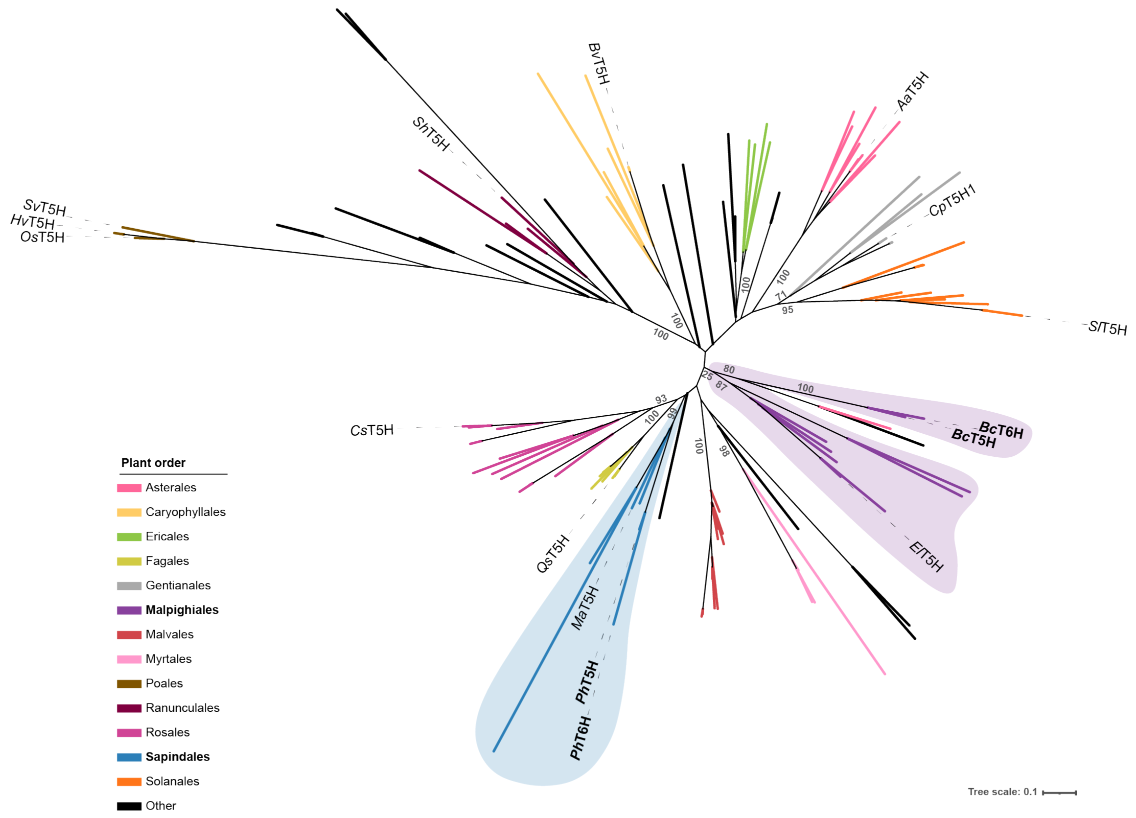

**SI Fig. 20. Maximum likelihood phylogenetic tree of tryptamine hydroxylases.** Labeled branches represent biochemically characterized enzymes. Branches are colored by plant phylogenetic order. Branch support for main nodes is indicated at the tree nodes by ultrafast bootstrap approximation (UFBoot) values using 1000 bootstrap resamplings. A complete description of sequences and parameters used can be found in the Methods section.

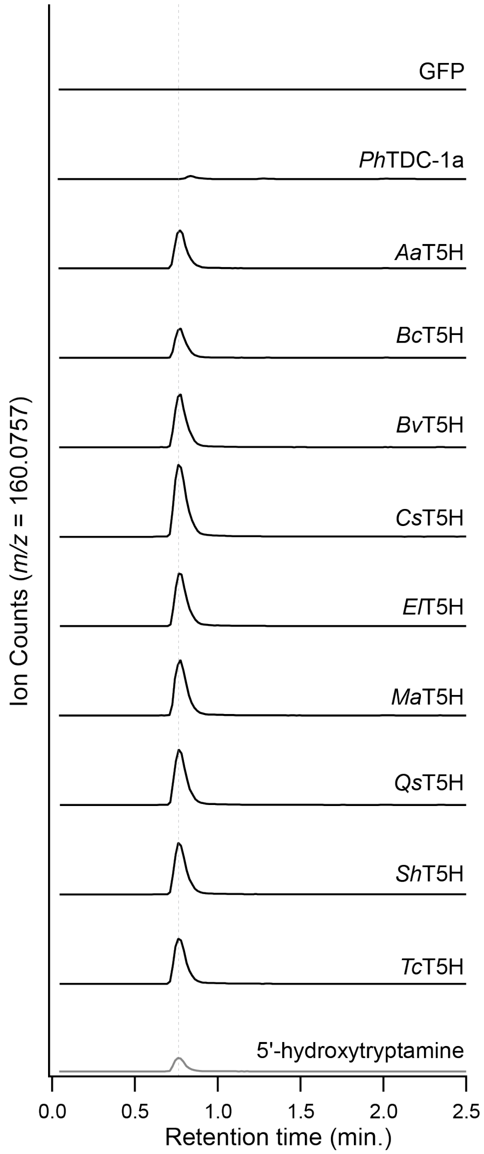

**SI Fig. 21. Characterization of putative tryptophan 5'-hydroxylases (T5Hs) from diverse plant orders.** T5H enzyme candidates were tested via transient expression in *N. benthamiana*. Each was co-expressed with *Ph*TDC-1a, which generates tryptamine (**1**) as an *in planta* substrate. The putative T5H enzymes are as follows: *Artemisia annua* (CYP71BG50; *Aa*T5H), *Beta vulgaris* (CYP71BG15; *Bv*T5H), *Cannabis sativa* (CYP71BG35b; *Cs*T5H), *Euphorbia lathyris* (CYP71BG51; *El*T5H), *Melia azedarach* (CYP71BG55; *Ma*T5H), *Quercus suber* (CYP71BG54; *Qs*T5H), *Sinopodophyllum hexandrum* (CYP71BG52; *Sh*T5H), *Theobroma cacao* (CYP71BG53; *Tc*T5H). Shown are LC-MS ion counts represent the major in-source fragment detected by LC-MS $[M\text{-}NH_3+H]^+$ = *m/z* 160.0757) rather than the parent ion ($[M+H]^+$ = *m/z* 177.1022).

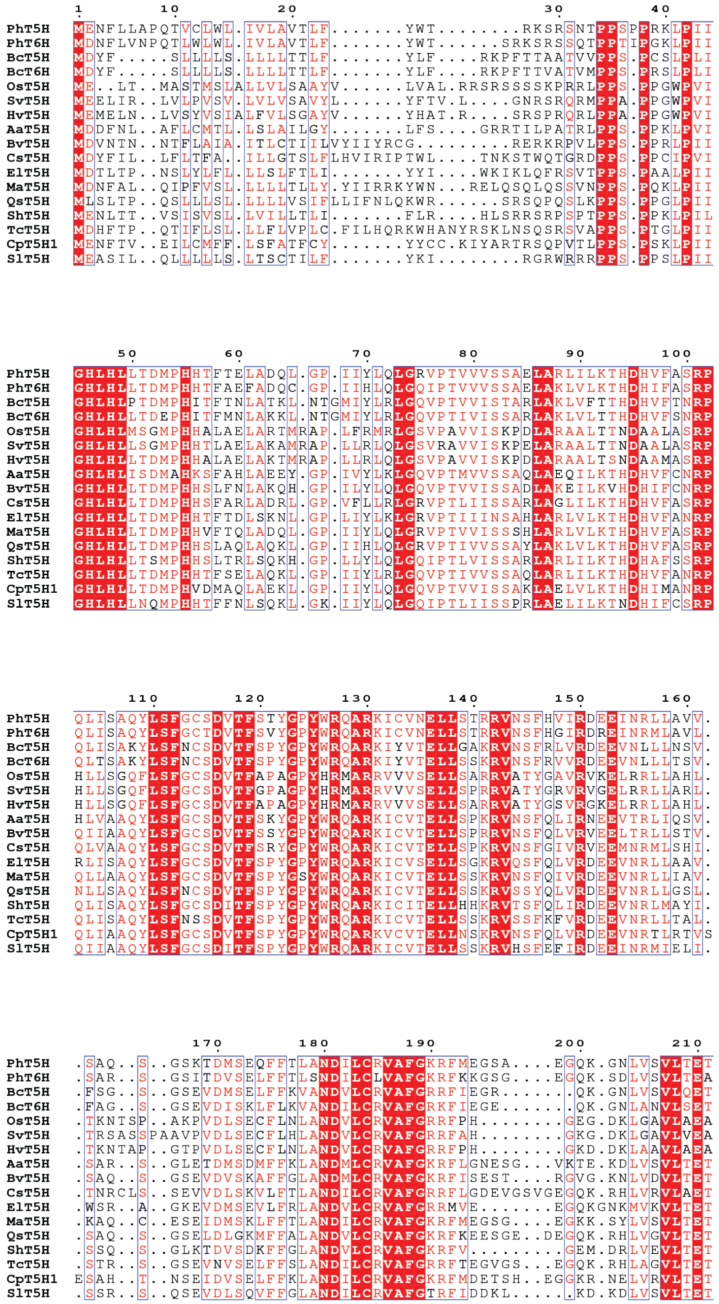

**SI Fig. 22. Protein multiple sequence alignment of characterized tryptamine hydroxylases.** Identical residues among all sequences analyzed are depicted in white font with red background. Residues with a similarity score above 70% are shown in red font and framed with blue frame. Blue triangles mark sites that are altered in either *Ph*T6H or *Bc*T6H compared to characterized T5H enzymes. These sites were selected for further mutagenesis studies.

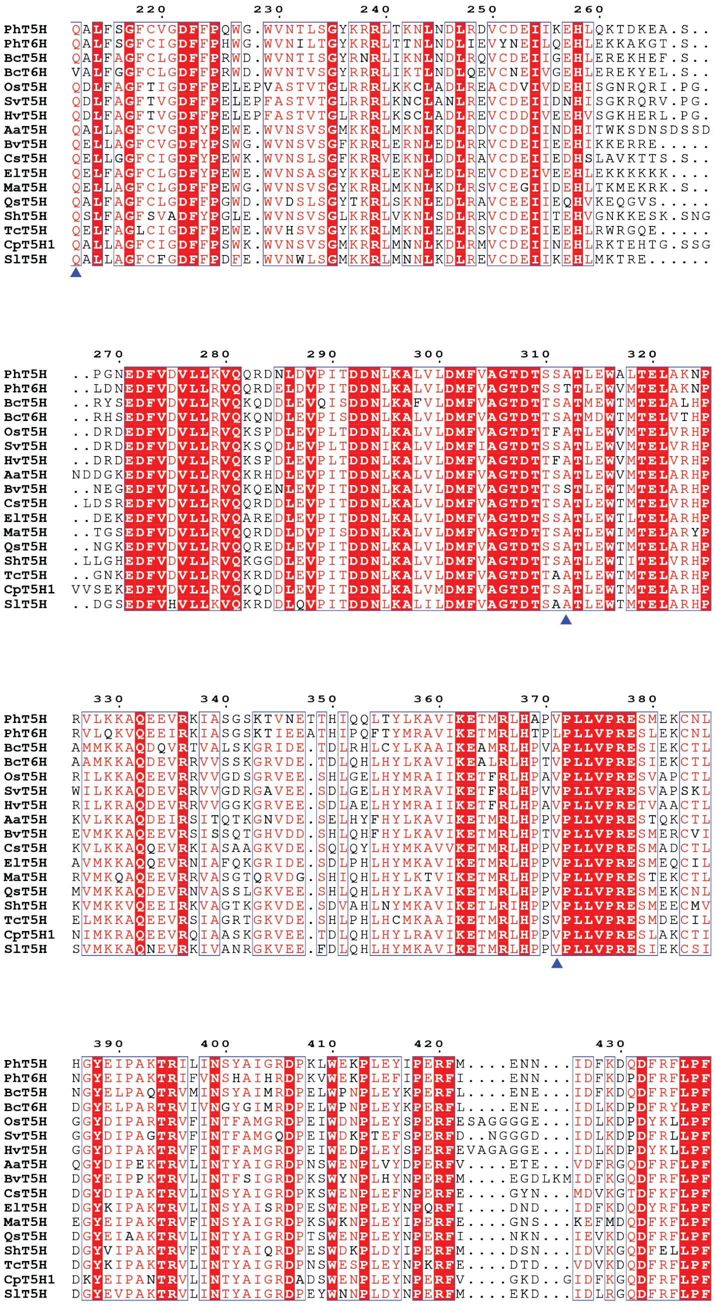

**SI Fig. 22. Continued (1)**

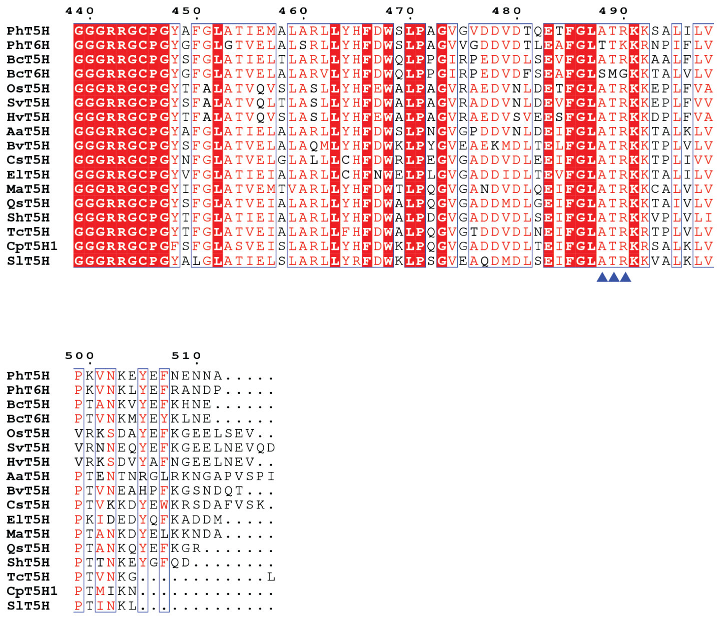

**SI Fig. 22. Continued (2)**

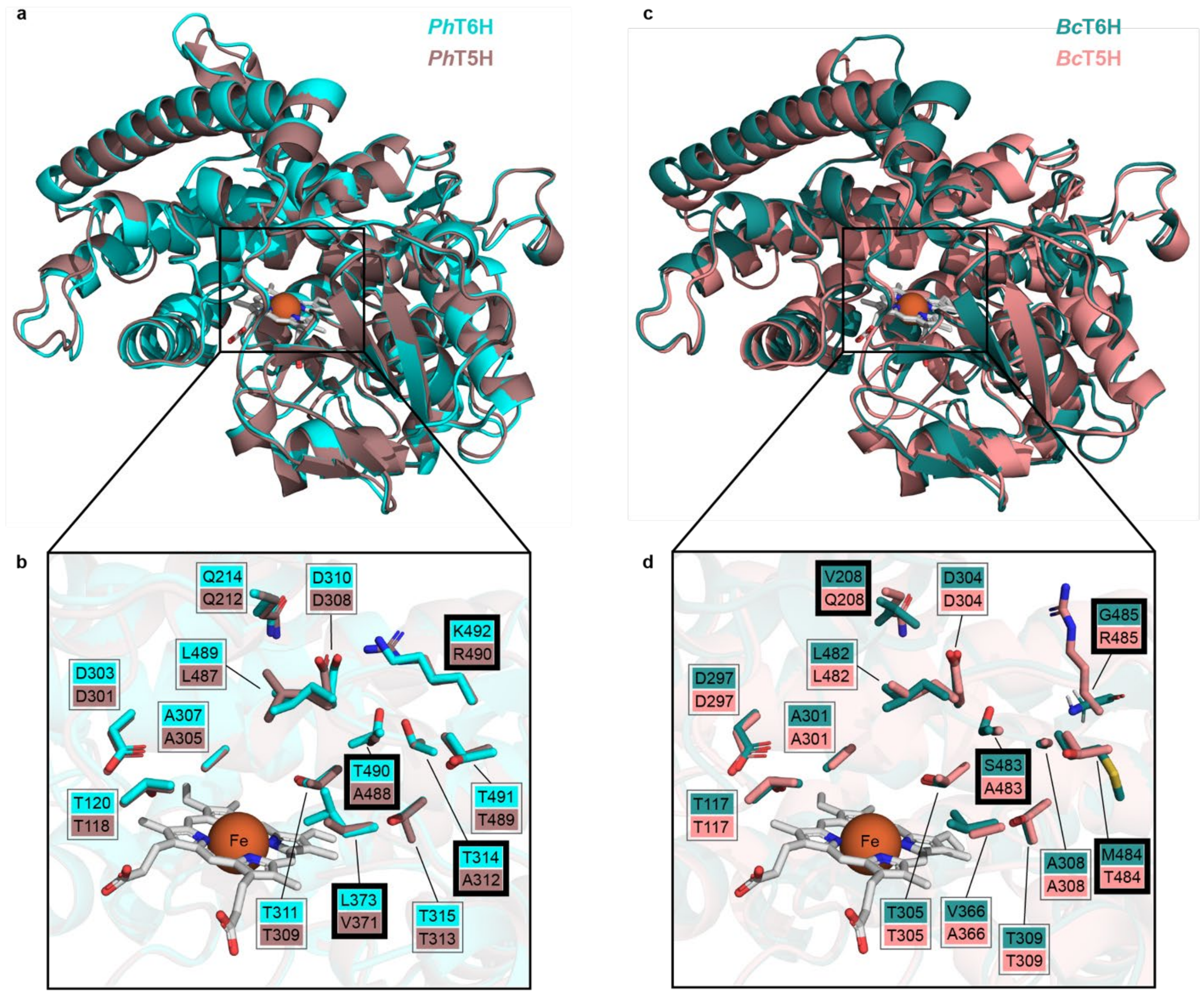

e

| Protein variant | Residue substitution |
|---|---|
| *Bc*T6H-1x | S483A |
| *Ph*T6H-1x | T490A |
| *Bc*T6H-4x | V208Q, S483A, M484T, G485R |
| *Ph*T6H-4x | T314A, L373V, T490A, K492R |

**SI Fig. 23. Comparison of predicted T6H and T5H structures from *P. harmala* and *B. caapi*.** AlphaFold3 was used to model the heme-bound structures of *Ph*T6H, *Ph*T5H, *Bc*T6H, and *Bc*T5H. Sequences were then aligned using the "super" function in PyMOL. N-terminal transmembrane domains were removed for visualization. **a**) Full alignment of *Ph*T6H and *Ph*T5H. **b)** Zoom in on the predicted active site for these enzymes and comparison of amino acid residues. Amino acids selected for mutagenesis are highlighted with bold boxes. **c**) Full alignment of *Bc*T6H and *Bc*T5H. **d)** Zoom in on the predicted active site for these enzymes and comparison of amino acid residues. Amino acids selected for mutagenesis are highlighted with bold boxes. **e)** T6H protein variants and their corresponding amino acid substitutions based on T5H conserved residues.

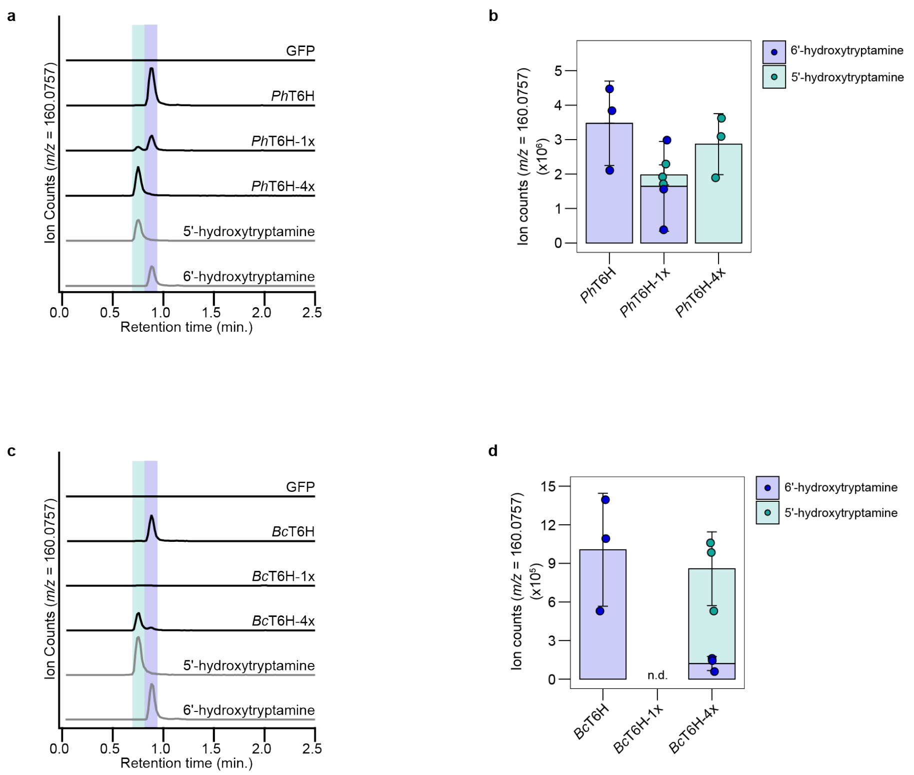

**SI Fig. 24. Characterization of mutagenized T6H enzymes from *P. harmala* and *B. caapi*.** Mutagenized T6H genes were transiently co-expressed with TDC genes in *N. benthamiana* and production of 5′-hydroxytryptamine and/or 6′-hydroxytryptamine (**2**) was quantified by LC-MS. **a**) EIC traces of *Ph*TDC-1a co-expressed with wild type *Ph*T6H, *Ph*T6H-1x (1 amino acid mutant), and *Ph*T6H-4x (4 amino acid mutant) compared to 5′-hydroxytryptamine and 6′-hydroxytryptamine standards ($[M\text{-}NH_3+H]^+$ = *m/z* 160.0757). **b**) Quantified EICs for 3 biological replicates of reactions described in panel (a). **c**) EIC traces of *Bc*TDC-1 co-expressed with wild type *Bc*T6H, *Bc*T6H-1x (1 amino acid mutant), and *Bc*T6H-4x (4 amino acid mutant) compared to 5′-hydroxytryptamine and 6′-hydroxytryptamine tryptamine (**2**) standards. **d**) Quantified EICs for 3 biological replicates of reactions described in panel (c). n.d. = not detected. In panels (b) and (d) dots represent biological replicates and bar graphs represent the mean ± s.d.

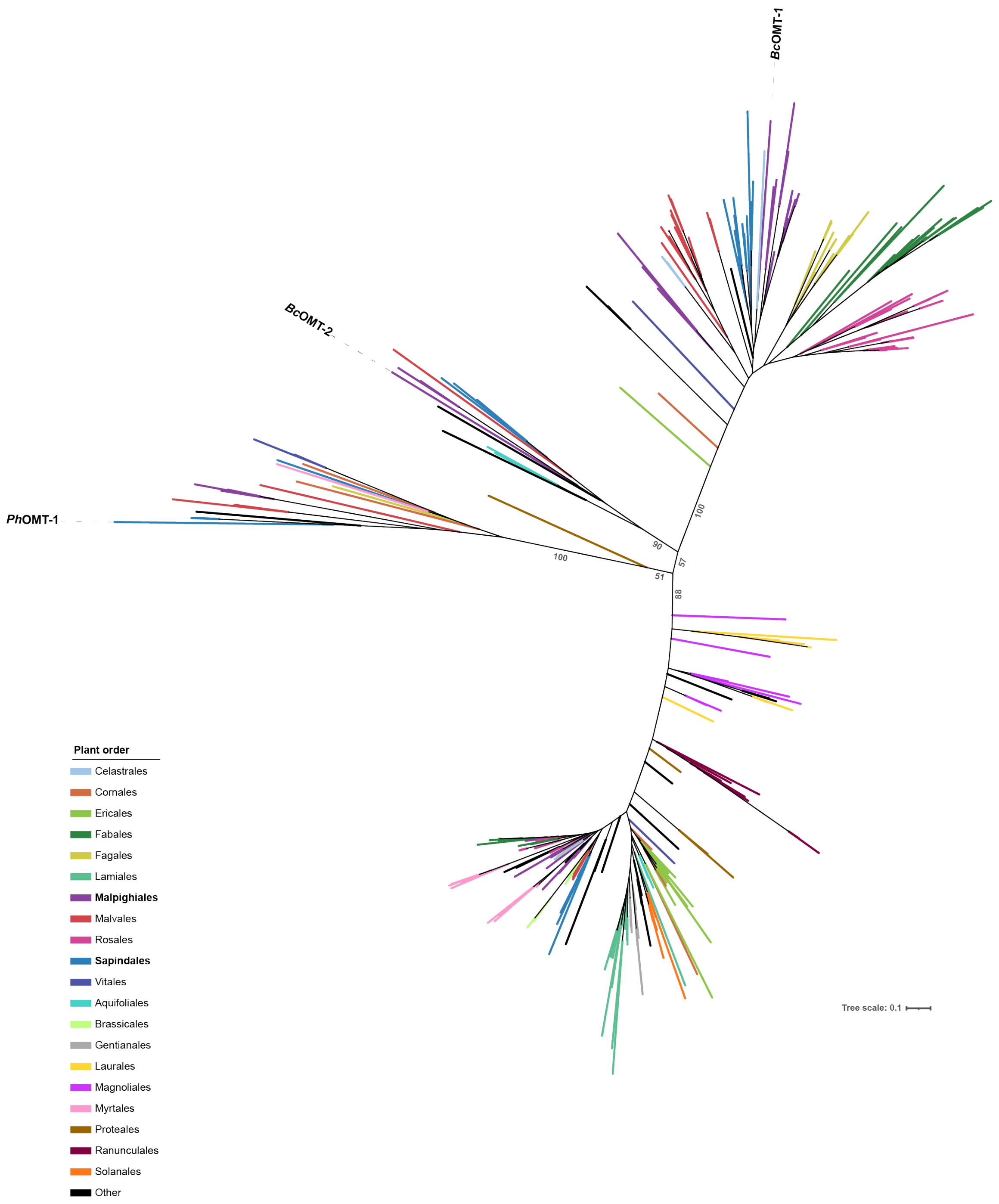

**SI Fig. 25. Maximum likelihood phylogenetic tree of *O*-methyltransferases (OMTs) evaluated in this study.** Labeled branches represent biochemically characterized enzymes. Branches are colored by plant phylogenetic order. Branch support for main nodes is indicated at the tree nodes by ultrafast bootstrap approximation (UFBoot) values using 1000 bootstrap resamplings. A complete description of sequences and parameters used can be found in the Methods section.

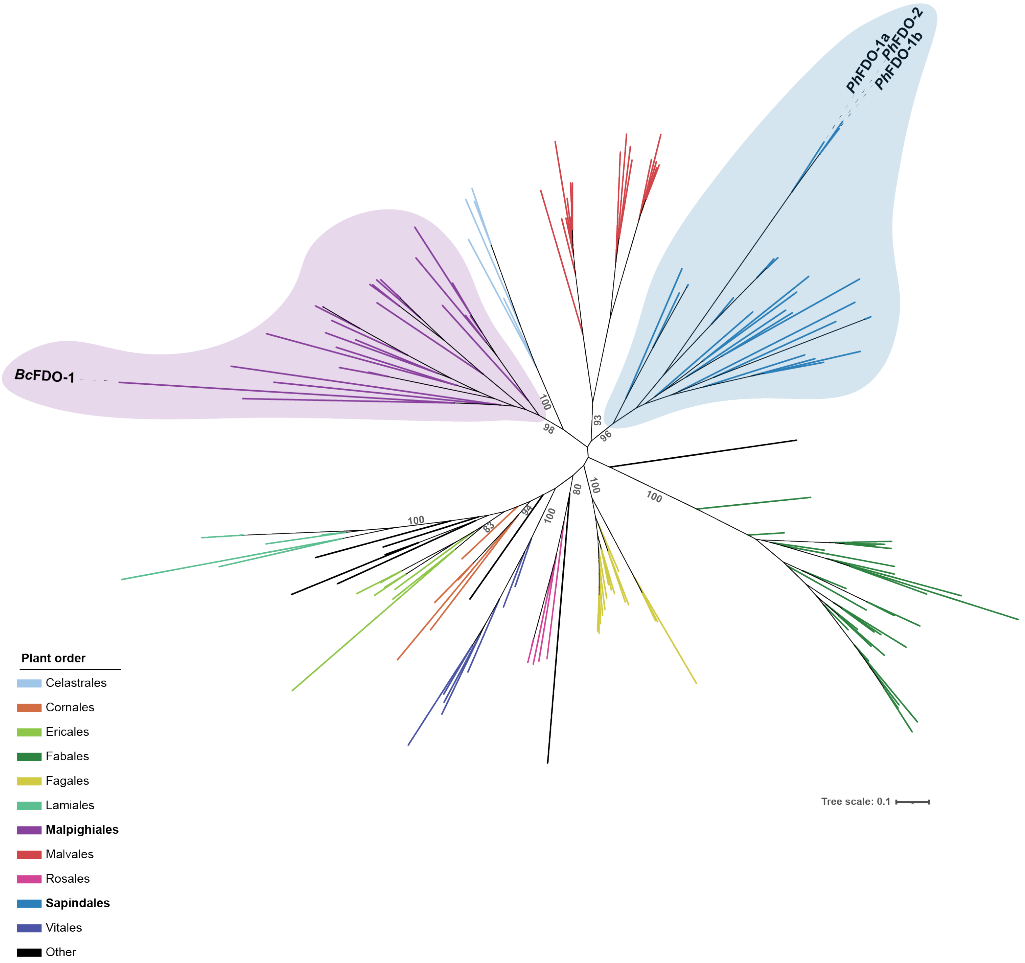

**SI Fig. 26. Maximum likelihood phylogenetic tree of flavin-dependent oxidases (FDOs) analyzed in this study.** Labeled branches represent biochemically characterized enzymes. Branches are colored by plant phylogenetic order. Branch support for main nodes is indicated at the tree nodes by ultrafast bootstrap approximation (UFBoot) values using 1000 bootstrap resamplings. A complete description of sequences and parameters used can be found in the Methods section.

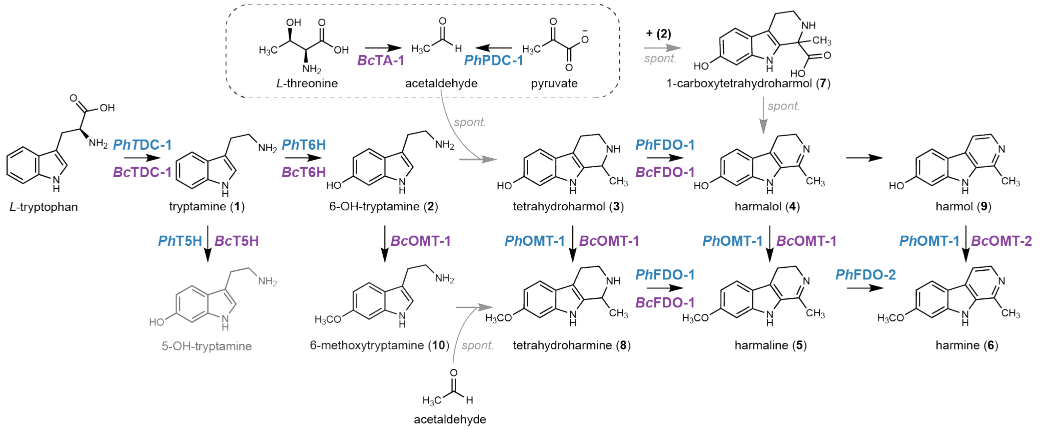

**SI Fig. 27. Summary of harmala alkaloid biosynthesis in *P. harmala* and *B. caapi*.** We note that 5-hydroxytryptamine is shown in gray as it is not on pathway to harmala alkaloids. spont. = spontaneous.

**Supplementary Methods**

Liquid chromatography parameters

| | |
|---|---|
| **Instrument** | Agilent 1290 Infinity II UHPLC system |
| **Column** | Agilent ZORBAX RRHD Eclipse Plus C18 (2.1 x 50 mm, 1.8 µm) |
| **Solvent A** | water with 0.1% (v/v) formic acid |
| **Solvent B** | acetonitrile with 0.1% (v/v) formic acid |
| **Flow rate** | 0.6 mL $min^{-1}$ |
| **Injection volume** | 1 µL |

| **% Solvent B** | **Time (min)** |
|---|---|
| 3 | 0.00 |
| 3 | 0.50 |
| 50 | 8.50 |
| 95 | 8.51 |
| 95 | 10.00 |
| 3 | 10.01 |
| 3 | 11.00 |

Mass spectrometry parameters

| **Instrument:** 6546 QTOF mass spectrometer | |
|---|---|
| **Mode** | Electrospray Ionization (ESI) positive |
| **Drying gas temperature** | 325˚C |
| **Drying gas flow rate** | 10 L $min^{-1}$ |
| **Nebulizer** | 35 psi |
| **Capillary voltage** | 4000 V |
| **Fragmentor** | 135 V |
| **Skimmer voltage** | 45 V |
| **Octopole 1 RF Vpp** | 750 V |
| **Acquisition time** | 1000 ms $spectrum^{-1}$ |
